# Rho/Rok-dependent regulation of actomyosin contractility at tricellular junctions restricts epithelial permeability in Drosophila

**DOI:** 10.1101/2024.10.04.616625

**Authors:** Thea Jacobs, Jone Isasti Sanchez, Steven Reger, Stefan Luschnig

**Affiliations:** Institute of Integrative Cell Biology and Physiology, Cells in Motion (CiM) Interfaculty Center, University of Münster, Röntgenstrasse 16, D-48149 Münster

**Keywords:** Cell vertex, tricellular junction, actin, myosin, Rho, oogenesis, follicle epithelium, epithelial permeability, *Drosophila*

## Abstract

Cell contacts in epithelia are remodeled to regulate paracellular permeability and to control passage of migrating cells, but how barrier function is modulated while preserving epithelial integrity is not clear. In the follicular epithelium of *Drosophila* ovaries, tricellular junctions (TCJs) open transiently in a process termed patency to allow passage of externally produced yolk proteins for uptake by the oocyte. Here we show that modulation of actomyosin contractility at cell vertices controls TCJ permeability. Before patency, circumferential actomyosin bundles are anchored at apical follicle cell vertices, where tension-sensing junctional proteins, Rho-associated kinase (Rok), and active Myosin II accumulate and maintain vertices closed. TCJ opening is initiated by redistribution of Myosin II from circumferential bundles to a medial pool, accompanied by decreasing tension on vertices. This transition requires activation of Cofilin-dependent F-actin disassembly by the phosphatase Slingshot and Myosin II inactivation by Myosin light chain phosphatase, and is counteracted by Rok. Accordingly, constitutive activation of Myosin or of Rho signaling prevent vertex opening, whereas reduced Myosin II or Rok activity cause excessive vertex opening. Thus, the opening of intercellular gaps in the follicular epithelium relies on relaxation of actomyosin contractility rather than active actomyosin-based pulling forces. Conversely, F-actin assembly is required for closing intercellular gaps after patency. Our findings are consistent with a force transduction model in which TCJ integrity is maintained by vertex-anchored contractile actomyosin. We propose that the cell-type-specific organization of actomyosin at cell vertices determines the mode of contractility-dependent regulation of epithelial permeability.

## Introduction

Intercellular contacts in epithelia are dynamically remodeled to enable cell rearrangements, paracellular transport, or passage of migrating cells ^1,2^. For instance, cell-cell contacts in the intestinal epithelium open transiently to enhance nutrient absorption after food intake ^3^. Paracellular extravasation of leukocytes from blood vessels requires the transient opening of endothelial cell junctions stimulated by pro-inflammatory cytokines that modulate vascular endothelial (VE)-Cadherin-based adhesion ^4^. Aberrant weakening or disassembly of intercellular junctions is a hallmark of infections and inflammatory conditions ^5,6^. Transient paracellular leaks also occur during cell division and cell rearrangements in epithelia and are normally rapidly repaired by localized activation of Rho GTPases to promote actin polymerization and actomyosin-mediated contraction of junctions ^7^. However, how epithelial cells modulate barrier function while preserving tissue integrity and undergoing dynamic shape changes during morphogenesis remains unclear.

Epithelial cells are connected by force-bearing adherens junctions (AJs) and by occluding junctions (tight junctions (TJs) in vertebrates and septate junctions (SJs) in invertebrates) that seal the intercellular space ^1^. The most abundant intercellular contacts in epithelia are bicellular junctions (BCJs) between two cells. However, at sites where three or more cells meet (cell vertices), specialized tricellular junctions (TCJs) composed of a distinct set of proteins mediate adhesion and paracellular sealing ^8^. In *Drosophila*, three TCJ-specific TM proteins, Anakonda (Aka; ^9^), Gliotactin (Gli; ^10^) and M6 ^11^ form the paracellular barrier along basolateral cell vertices, whereas the immunoglobulin domain TM protein Sidekick (Sdk) localizes at apical tricellular AJs (tAJs) and is required for efficient cell rearrangements ^12–15^. Vertebrate epithelia display tricellular tight junctions (tTJs; ^16^) that differ structurally from invertebrate TCJs and comprise TM proteins of the Angulin/Lipolysis-stimulated Lipoprotein Receptor (LSR) family ^17^ and the MARVEL family TM protein Tricellulin ^18^.

Cell vertices are key points that sense and integrate forces in epithelial and endothelial tissues ^19^. They are dynamically remodeled during cell rearrangements ^20^ and play fundamental roles in cytoskeletal organization, mitotic spindle orientation, and epithelial barrier function ^21^. Moreover, TCJs provide preferred routes for leukocyte extravasation from blood vessels ^22^ and are exploited by bacterial pathogens for breaching epithelial barriers ^23^. However, despite their crucial functions, how TCJs are assembled and remodeled is only beginning to be understood.

Studies in *Drosophila* have revealed important insights into the molecular mechanisms underlying epithelial cell polarization and intercellular junction formation ^1,24^. *Drosophila* ovaries contain chains of developing follicles, each of which comprises 16 germline cells encased by an epithelial monolayer of somatic follicle cells (FCs; Fig. 1A). TCJs in the follicular epithelium overlying the oocyte open transiently during vitellogenesis to generate intercellular channels that allow passage of externally produced yolk proteins for uptake by the oocyte ^25,26^. This process, referred to as follicular patency ^27,28^, provides an accessible model for investigating the mechanisms of TCJ remodeling. The opening of TCJs is initiated by the sequential removal of adhesion proteins (E-Cad, N-Cad, NCAM/Fasciclin 2, Sdk) from FC vertices and by actomyosin remodeling ^29,30^. Decreasing levels of non-muscle Myosin II (MyoII) and of MyoII regulators in FCs at the onset of patency suggested that downregulation of MyoII contractility is necessary for opening of vertices ^29^. However, how actomyosin is organized and remodeled at cell vertices and how contractility is modulated to control TCJ permeability was not clear.

**Figure 1.**
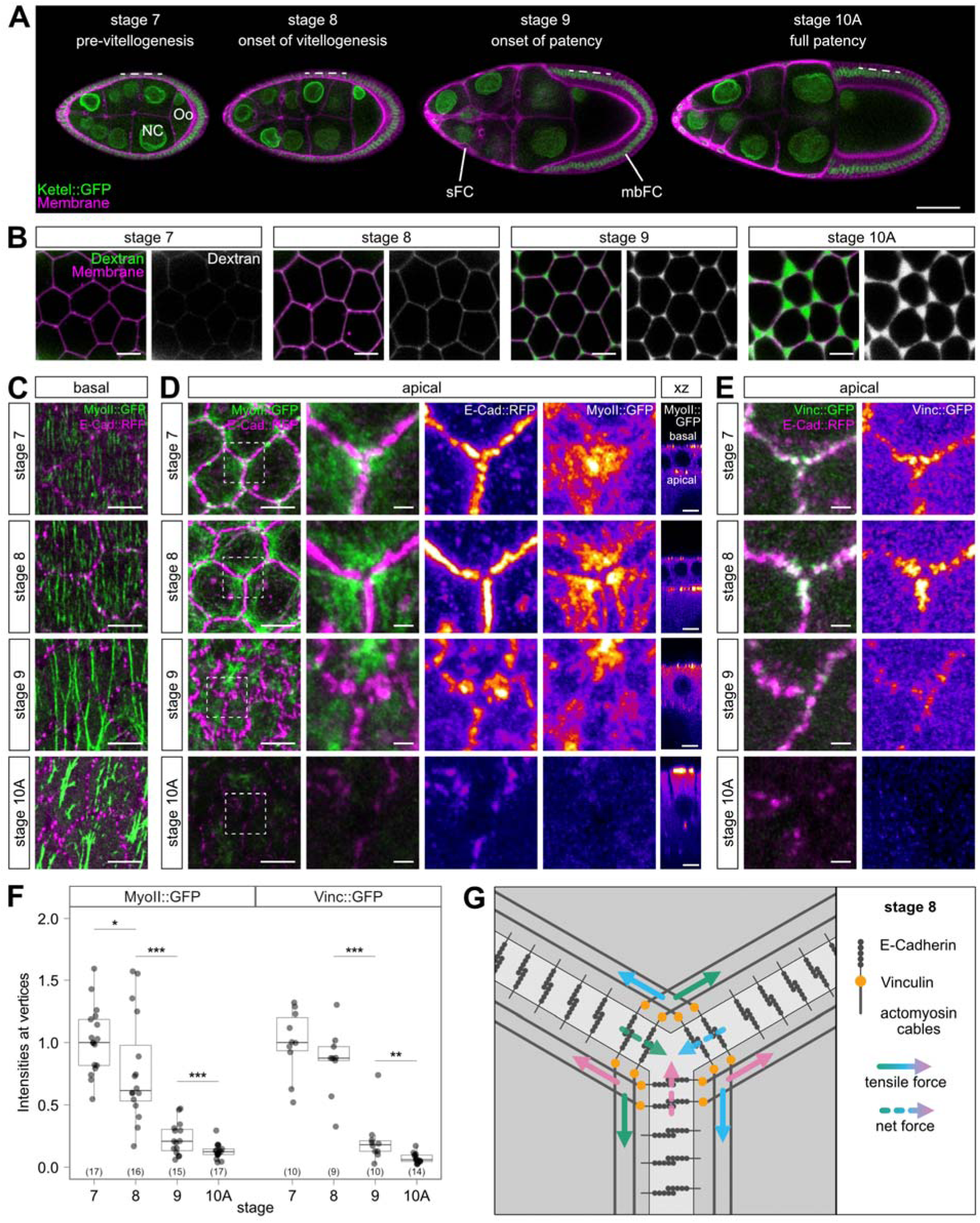
Reorganization of actomyosin at follicle cell vertices precedes vertex opening. **(A)** Confocal sections of living follicles before (stage 7) and during (stages 8, 9, 10A) vitellogenesis. Fs(2)Ket::GFP (green) labels nuclei, CellMask (magenta) labels plasma membranes. Anterior is to the left, dorsal is up. NC, nurse cells; Oo, oocyte; BC, border cells; sFC, stretch follicle cells; mbFC, main body follicle cells. **(B)** Basolateral sections of mbFCs acquired at 25% FC height (dashed lines in A). Dextran (green) marks intercellular spaces, CellMask (magenta) labels plasma membranes. Single-channel views show dextran (grey). **(C-D)** Basal (C) and apical (D) sections of mbFCs expressing E-Cad::RFP (magenta) and MyoII::GFP (green). Close-ups (D) show single vertices (rectangles in overview). Note that MyoII (intensities color-coded using heatmap in (D)) accumulates near apical vertices at stage 7 and 8 and redistributes to the medial region at stage 9 (D). Cross-sectional (xz) view (D) shows that apical, but not basal, MyoII declines by stage 9. **(E)** Apical sections of FC vertices in follicles expressing E-Cad::RFP (magenta) and Vinc::GFP (green). Note that Vinc::GFP (color-coded using heatmap in single-channel view) accumulates near vertices at stage 7 and 8 and declines by stage 9. **(F)** Quantification of MyoII::GFP and Vinc::GFP signals (normalized) at apical vertices. Box plots show maximum and minimum observations, upper and lower quartile, and median (horizontal line). Number of follicles (n) analyzed is indicated. P-values (Pairwise Wilcoxon Rank Sum Test): * p ≤ 0.05, ** p ≤ 0.01, *** p ≤ 0.001, **** p ≤ 0.0001. **(G)** Model of force transmission at apical FC vertices at stage 8 (modified from ^19^). Scale bars: (A), 50 µm; (B-D), 5 µm, close-ups 1 µm; (E) 1 µm. See also Figure S1 and Video S1.

We show here that vertex-anchored actomyosin generates tensile forces that keep TCJs closed before patency. This depends on Rho-associated kinase (Rok), a main effector of Rho GTPase ^31,32^. Rok stimulates actomyosin contractility in mammalian cells by phosphorylating MyoII regulatory light chain (MRLC), thus activating MyoII, and by inhibitory phosphorylation of Myosin light-chain phosphatase (MLCP), which normally inactivates MyoII ^31,33^. In parallel, Rok regulates F-actin dynamics by activating LIM-domain kinase 1 (LIMK1), which in turn inactivates the F-actin-severing protein Cofilin ^34–36^. We found that in *Drosophila* FCs, Rok is required to maintain active MyoII, but not for regulating F-actin dynamics via the LIMK1-Cofilin pathway. At the onset of patency, downregulation of Rok results in MyoII inactivation and decreased tension acting on vertices. In parallel, the protein phosphatase Slingshot (Ssh; ^37^) activates Cofilin-dependent F-actin disassembly. This work reveals how actomyosin contractility at TCJs is modulated to control epithelial permeability.

## Results

### Reorganization of apical actomyosin precedes vertex opening

TCJs in the main body follicle epithelium overlying the oocyte open during vitellogenesis (stages 8-10; Fig. 1A; ^25^) to generate patent intercellular channels, which are visualized by incubating follicles in fluorescent dextran (Fig. 1B; ^29^). Intercellular gaps first appear at stage 9, reach maximal width at stage 10A, and close at stage 11 (Fig. 1B; ^29,30^).

To understand the mechanism of vertex opening, we investigated the distribution of actomyosin in main body FCs at the onset of patency. We visualized E-Cadherin (E-Cad::RFP), MyoII (MyoII::GFP) and the mechanosensitive protein Vinculin (Vinc::GFP) in living follicles before (stages 7 and 8) and during early (stage 9) and full patency (stage 10A; Fig. 1C-E). On the basal side of FCs, MyoII was enriched in planar polarized stress fibers throughout the stages analyzed (Fig. 1C; ^38,39^). By contrast, on the apical side of FCs in pre-patent follicles (stages 7 and 8), MyoII (Fig. 1D) and actin filaments (Lifeact::mBaoJin; Fig. S1A,B) aligned at the cell perimeter, with prominent accumulations at TCJs. MyoII bundles appeared to connect to vertices, where Vinc was enriched overlapping with E-Cad (Fig. 1D,E), suggesting that actomyosin bundles exert tension on AJs at cell vertices. While E-Cad was present in continuous belt-like AJs at stage 7, E-Cad was removed from vertices at stage 8 (Fig. 1D). At onset of patency (stage 9), MyoII receded from vertices and redistributed from circumferential bundles to a medial pool (Fig. 1D; ^40^). Apical MyoII levels further decreased by stage 10A, while basal MyoII-containing stress fibers were maintained (Fig. 1D). Concurrently, Vinc accumulations at apical vertices decreased by 74.7% (n(st8)=9, n(st9)=10, p=0.00026) from stage 8 to stage 9 and disappeared by stage 10A (Fig. 1E). Similarly, the force-sensing LIM-domain proteins Ajuba (GFP::Jub) and Zyxin (Zyx::GFP) ^41^ accumulated in distinct puncta adjacent to apical vertices at stage 8 and disappeared by stage 9 (Fig. S1C-F). These findings suggest that apical FC vertices are under tension in pre-patent follicles and that tension on vertices decreases at the onset of patency.

To explain these results, we considered a force transduction model ^19,42–44^ in which contractile actomyosin bundles make end-on connections to AJs at vertices (Fig. 1G). The combined forces acting on these AJs are directed towards the center and tighten the vertex. Conversely, reduced actomyosin-based tension at stage 9 is predicted to release tension and thereby allow opening of vertices for patency.

### Tension on vertices decreases at the onset of patency

To test this model, we asked whether vertices in pre-patent FCs are indeed under tension. We laser-ablated apical FC vertices (marked by E-Cad::GFP) at stage 7, 8 or 9, and analyzed displacement of neighboring vertices as a measure for tension acting on them (Fig. S2A,B; Video S1). This revealed rapid displacement in stage 7 (0.0114 +/- 0.0063 µm/s) and stage 8 (0.0097 +/- 0.0048 µm/s) follicles. By contrast, displacement was strongly reduced (0.0038 +/- 0.0023 µm/s) at stage 9 (Fig. S2B-D), consistent with results from ablations of bicellular junctions ^40^. These findings, along with the behavior of force-sensing proteins (Fig. S1), indicate that FC vertices are under tension in pre-patent follicles and suggest that tension on vertices decreases before the onset of patency.

### Rho-associated kinase localizes at follicle cell vertices and limits vertex opening

To understand how tension on vertices is generated, we analyzed the distribution of Rok, a key regulator of actomyosin contractility ^32,33^. We found that an mNeonGreen::Rok fusion protein (mNG::Rok; ^45^) accumulates at apical cell boundaries, with prominent puncta abutting FC vertices in pre-patent follicles (Fig. 2A). Like Rok, active MyoII, detected by an antibody against MRLC phosphorylated on Serine 21 (p-MyoII), accumulated near apical vertices in pre-patent follicles (Fig. 2B). Rok puncta disappeared by the onset of patency at stage 9 (Fig. 2A,E), resembling the behavior of tension-sensing proteins Vinc, Jub, and Zyx (Fig. 1E, Fig. S1). Along with mNG::Rok, p-MyoII accumulations receded from vertices towards the medial region at stage 9, suggesting that Rok maintains a pool of active MyoII at vertices. Supporting this idea, RNAi-mediated depletion of Rok in FC clones led to a substantial reduction (57.9% of control (st. 8); n=11, p=0.000881) of p-MyoII signals at vertices (Fig. 2C,F). Rok-depleted cells showed enlarged apical outlines, indicating reduced apical constriction, as reported previously ^40,46^. Strikingly, Rok-depleted FC clones also displayed enlarged dextran-filled intercellular gaps (quantified as the fraction of total cell area, referred to as patency index ^29^; Fig. 2D,G). Intercellular gaps in Rok-depleted tissue occasionally appeared before the onset of patency in neighboring control tissue (Fig. 2D), but were generally not observed before stage 8. Consistent with this finding, E-Cad was present at vertices of Rok-depleted cells comparable to wild-type cells at stage 8 (Fig. 2C,F). Together, these results suggest that Rok-dependent MyoII activity is required to limit the extent of vertex opening after removal of adhesion proteins from vertices permits the initiation of patency.

**Figure 2.**
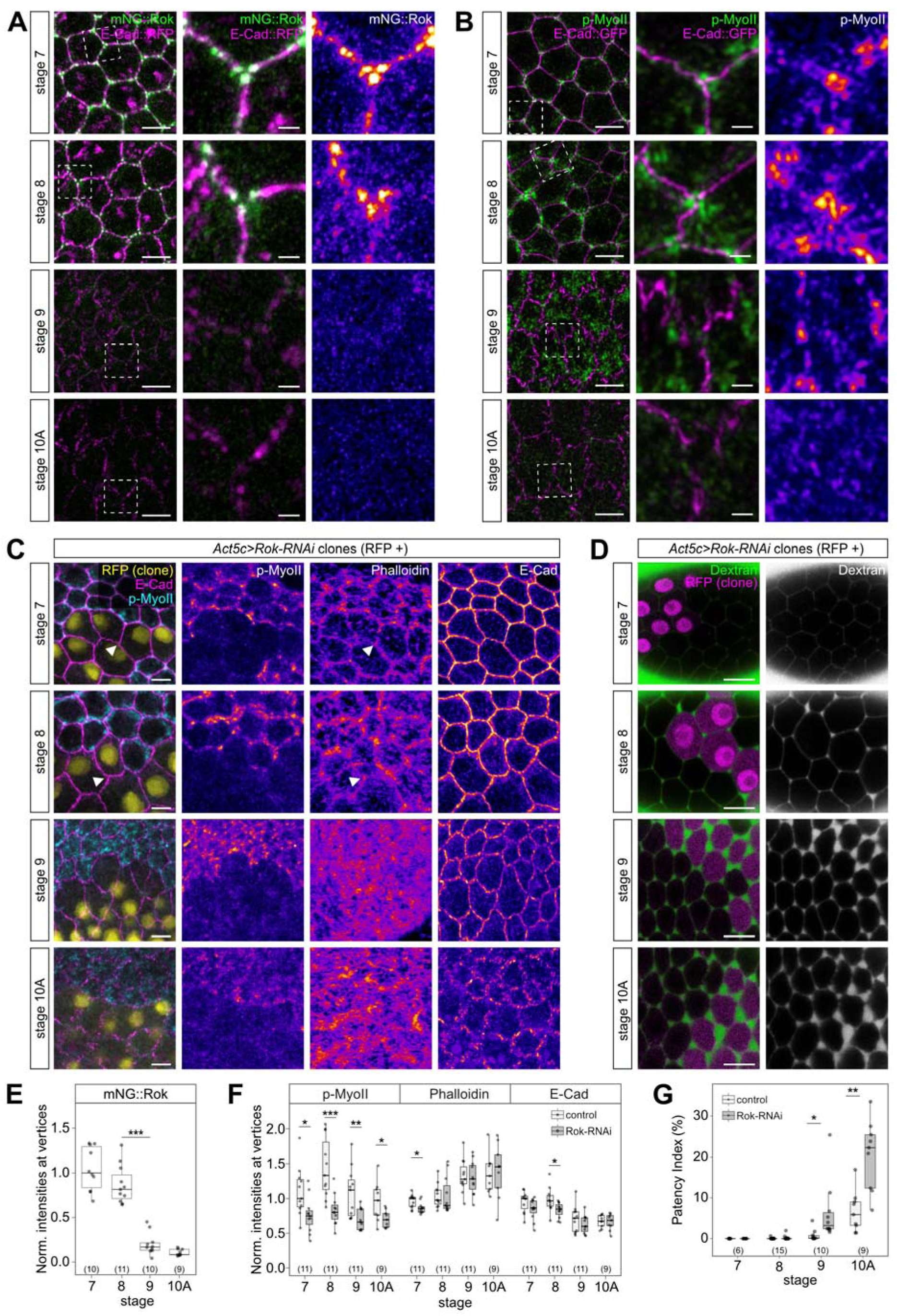
Rho-associated kinase localizes at follicle cell vertices and limits vertex opening. **(A)** Apical sections of mbFCs expressing E-Cad::RFP (magenta) and mNeonGreen::Rok (mNG::Rok; green). Close-up views show single vertices. Note that mNG::Rok (intensities color-coded using heatmap in single-channel view) accumulates near vertices in pre-patent follicles (stage 7, 8), but not in later stages. **(B)** Apical sections of follicles expressing E-Cad::GFP (magenta) stained for p-MyoII (green; heatmap in single-channel view). Close-ups show single vertices. Note that p-MyoII redistributes from vertices to the medial region by stage 9. **(C)** Apical sections of follicles expressing *Rok* dsRNA in RFP-nls-marked clones (yellow) stained for p-MyoII (cyan), E-Cad (magenta) and F-actin (Phalloidin; not shown in merge). Note reduced p-MyoII, but not F-actin signals in Rok-depleted cells. Arrowheads indicate F-actin accumulation at vertices in Rok-depleted cells. **(D)** Basolateral sections of follicles expressing *Rok* dsRNA in RFP-nls-marked clones (magenta). Dextran (green) marks intercellular spaces. Note enlarged gaps in Rok-depleted cells compared to control cells at stage 9 and stage 10A. **(E)** Quantification of mNeonGreen::Rok signals (normalized) at apical vertices. Number of follicles (n) analyzed is indicated. P-values (Pairwise Wilcoxon Rank Sum Test): * p ≤ 0.05, ** p ≤ 0.01, *** p ≤ 0.001, **** p ≤ 0.0001. **(F)** Quantification of p-MyoII, Phalloidin and E-Cad signals (normalized) at apical vertices in *Rok* RNAi and control cells. Number of follicles (n) analyzed is indicated. P-values (Welch’s two-sample t-test): * p ≤ 0.05, ** p ≤ 0.01, *** p ≤ 0.001, **** p ≤ 0.0001. **(G)** Quantification of patency index in *Rok* RNAi and control cells. Number of follicles (n) analyzed is indicated. P-values (Welch’s two-sample t-test): * p ≤ 0.05, ** p ≤ 0.01, *** p ≤ 0.001, **** p ≤ 0.0001. Scale bars: (A-B), 5 µm, close-ups 1 µm; (C) 5 µm; (D) 10 µm. See also Figures S2 and S3.

### Downregulation of Myosin II activity by protein phosphatase 1 is required for patency

We asked next how Rok regulates MyoII activity in FCs. Rok can activate Myosin by phosphorylating MRLC and by inhibiting Myosin light chain phosphatase (MLCP, also known as protein phosphatase 1 (PP1), which inactivates MyoII by dephosphorylating MRLC (Fig. 3A) ^33^. We found that loss of the PP1 catalytic subunit, PP1β, encoded by *flapwing (flw*; ^47,48^), in *flw^FP41^* FC clones caused excessive MyoII activation (indicated by elevated p-MyoII levels) in pre-patent follicles (Fig. 3B-D). MyoII hyperactivation, accompanied by accumulation of F-actin, in *flw^FP41^* clones persisted into stage 10A, when apical p-MyoII levels are low in wild-type FCs (Fig. 3B-D, Fig. 3F), suggesting that PP1β normally inactivates apical MyoII at the onset of patency. *flw^FP41^* cells showed smaller apical areas compared to control cells (Fig. 3F), presumably due to increased apical constriction upon MyoII overactivation. Strikingly, FC vertices in *flw^FP41^* clones failed to open when surrounding wild-type epithelium was fully patent at stage 10A (Fig. 3G,H), indicating that MyoII inactivation by PP1β is required for FCs to enter patency. Conversely, interfering with MyoII activity by expressing in FCs a dominant-negative form of Myosin II heavy chain lacking the actin-binding head domain (MHC^DN^::YFP; ^49^) caused reduced MyoII phosphorylation (Fig. 3K) and excessive vertex opening (Fig. 3I,J). Together, these findings indicate that Rok-dependent MyoII contractility acts to keep FC vertices closed until stage 9, while subsequent downregulation of MyoII activity by MLCP is necessary to allow vertex opening, the extent of which is determined by the degree of MyoII activity.

**Figure 3.**
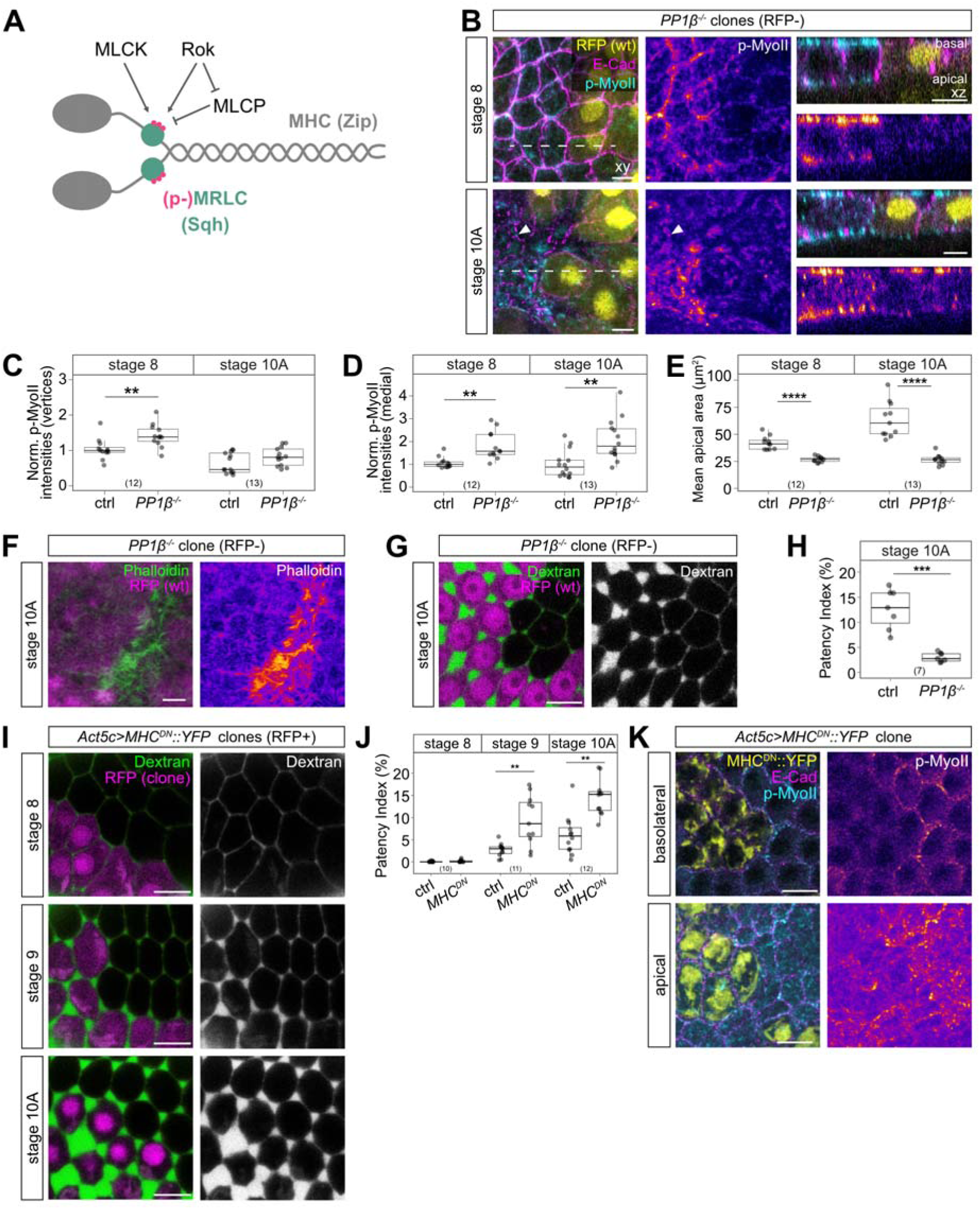
Downregulation of Myosin II activity by protein phosphatase 1 is required for patency. **(A)** Scheme of non-muscle Myosin II with Myosin heavy chain (MHC/Zip), Myosin regulatory light chain (MRLC/Sqh), and its regulators Myosin light chain kinase (MLCK), Myosin light-chain phosphatase (MLCP), and Rok. **(B)** Apical (xy, left) and orthogonal (xz, right) sections of follicle epithelium carrying *PP1β* (*flw^FP41^*) clones marked by absence of RFP (yellow), stained for E-Cad (magenta) and p-MyoII (cyan). Dashed lines indicate positions of orthogonal sections. Note increased p-MyoII levels (heatmap) on apical and basal sides in *PP1β* mutant cells. Arrowhead indicates p-MyoII accumulation at vertex in *PP1β* mutant cells. **(C)** Quantification of p-MyoII signals (normalized) at apical vertices in *PP1β* and control cells. Number of follicles (n) analyzed is indicated. P-values (Welch’s two-sample t-test): * p ≤ 0.05, ** p ≤ 0.01, *** p ≤ 0.001, **** p ≤ 0.0001. **(D)** Quantification of p-MyoII signals (normalized) at apical medial region in *PP1β* and control cells. Number of follicles (n) analyzed is indicated. P-values (Welch’s two-sample t-test): * p ≤ 0.05, ** p ≤ 0.01, *** p ≤ 0.001, **** p ≤ 0.0001. **(E)** Quantification of apical area in *PP1β* and control cells. Number of follicles (n) analyzed is indicated. P-values (Welch’s two-sample t-test): * p ≤ 0.05, ** p ≤ 0.01, *** p ≤ 0.001, **** p ≤ 0.0001. **(F)** Apical section of follicle epithelium carrying *PP1β* (*flw^FP41^*) clone marked by absence of RFP (magenta) reveals F-actin (Phalloidin; green; heatmap in single-channel view) accumulation in *PP1β* mutant tissue. **(G)** Basolateral section of follicle carrying *PP1β* clone marked by absence of RFP (magenta). Dextran (green) marks intercellular spaces. **(H)** Quantification of patency index of *PP1β* and control cells. Number of follicles (n) analyzed is indicated. P-values (Welch’s two-sample t-test): * p ≤ 0.05, ** p ≤ 0.01, *** p ≤ 0.001, **** p ≤ 0.0001. **(I)** Basolateral sections of mbFCs expressing MHC^DN^::YFP in RFP-nls-marked clones (magenta). Dextran (green) marks intercellular spaces. Note enlarged intercellular gaps in MHC^DN^::YFP-expressing tissue at stage 9 and 10A. **(J)** Patency index of control and MHC^DN^::YFP-expressing clones. Number of follicles (n) analyzed is indicated. P-values (Welch’s two-sample t-test): * p ≤ 0.05, ** p ≤ 0.01, *** p ≤ 0.001, **** p ≤ 0.0001. **(K)** Stage 10A follicle expressing MHC^DN^::YFP (yellow) in clones, stained for E-Cad (magenta) and p-MyoII (cyan). Note that MHC^DN^::YFP-expressing cells show reduced p-MyoII levels. Scale bars: (B, F) 5 µm; (G, I, K) 10 µm. See also Figure S3.

### Rok controls patency by regulating Myosin II activity

Besides promoting MyoII contractility, Rok stimulates F-actin assembly in mammalian cells by activating LIMK1, which in turn inhibits the F-actin-severing activity of Cofilin ^34–36^. To test whether Rok-dependent F-actin regulation is involved in vertex remodeling, we analyzed the effects of *Rok* RNAi on F-actin in FCs. Surprisingly, while *Rok* RNAi led to substantially reduced apical p-MyoII signals, there was only a minor reduction of F-actin in Rok-depleted cells at stage 7 and no significant effect on the distribution or levels of F-actin at later stages (Fig. 2C,F). These findings suggest that in FCs Rok acts mainly by controlling MyoII activity, whereas Rok-dependent regulation of F-actin dynamics via LIMK1 appears to play no significant role. Consistent with this notion, loss of LIMK1 in homozygous viable *LIMK1*^2^ null mutants ^50^ had no detectable effect on F-actin organization in FCs or on patency (Fig. S3).

### Vertex-associated F-actin prevents TCJ opening

F-actin was enriched at apical FC vertices in pre-patent follicles (stages 7, 8) and receded from vertices by stage 9 (Fig. S1A,B), suggesting that initiation of patency involves F-actin remodeling. To test this idea, we used mutations or RNAi to probe the roles of F-actin-regulators. However, in many cases this led to pleiotropic effects on epithelial organization or cell survival (not shown), thus precluding specific functions in patency. Therefore, to deplete F-actin in a more acute fashion, we treated cultured follicles with Latrunculin A (LatA), which prevents actin assembly by sequestering actin monomers and by accelerating F-actin depolymerization ^51^. In follicles treated with LatA for 45 minutes, apical F-actin was strongly reduced compared to controls (Fig. 4A,B) and E-Cad-labelled AJs were interrupted at vertices (Fig. 4D). Interestingly, LatA treatment led to enlarged intercellular gaps on the apical, but not basolateral side (Fig. 4C-E). An increase in apical gap size after LatA treatment was detectable (albeit with low significance; p ≤ 0.05) already at stage 8 and robustly from stage 9 onwards (Fig. 4D,E). These findings suggest that an apical pool of LatA-sensitive F-actin is required to maintain E-Cad at vertices and to prevent precocious vertex opening.

**Figure 4.**
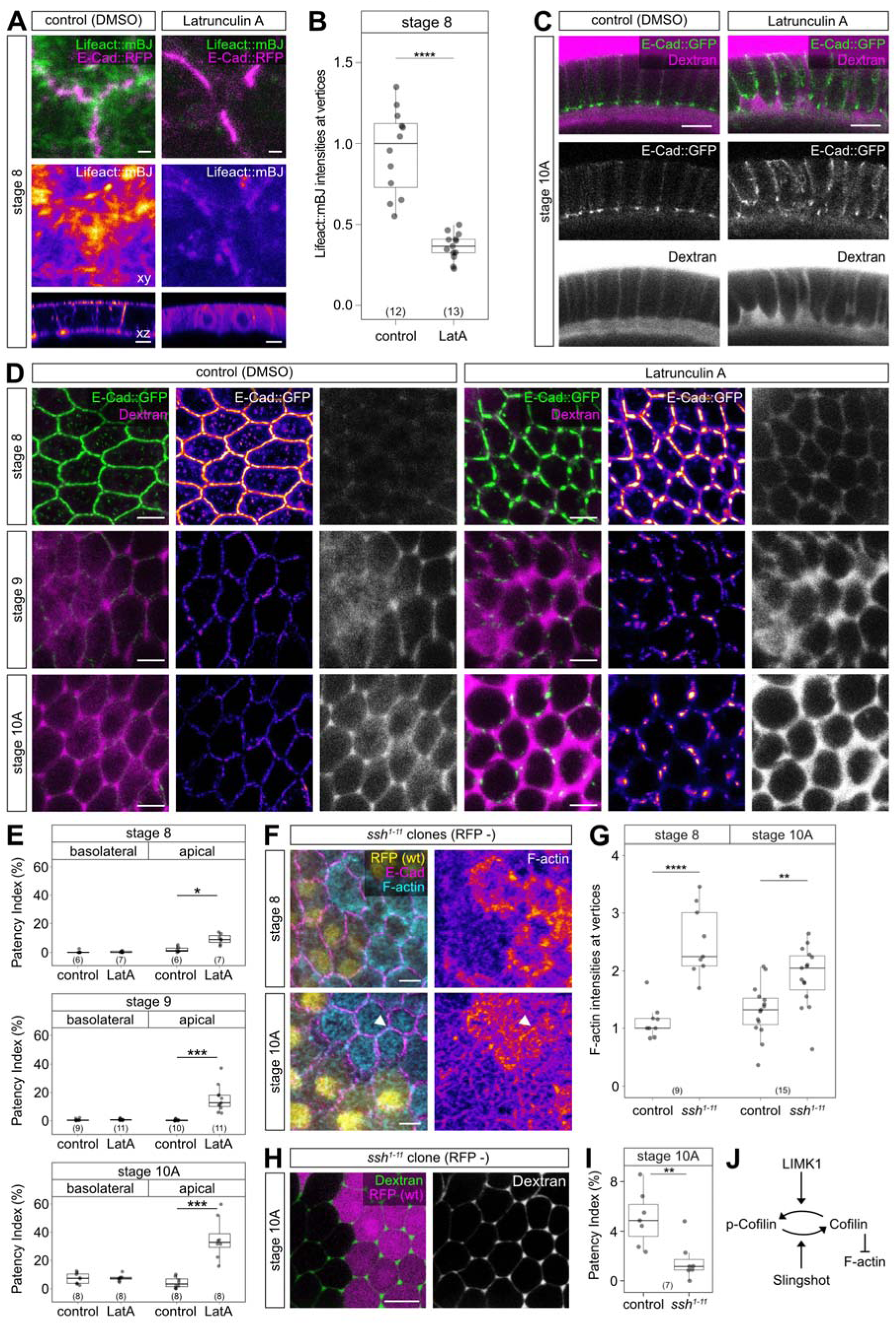
F-actin dynamics regulates vertex opening. **(A)** Apical (xy) or orthogonal (xz) sections of stage 8 follicles expressing E-Cad::RFP (magenta) and Lifeact::mBaoJin (green; driven by *tj*-Gal4), treated with DMSO (1%, control) or Latrunculin A (LatA; 2 µM). Note reduced Lifeact::mBaoJin intensities after LatA treatment. **(B)** Quantification of Lifeact::mBaoJin signals (normalized) at apical vertices of stage 8 follicles treated with DMSO (control) or LatA. Number of follicles (n) analyzed is indicated. P-values (Welch’s two-sample t-test): * p ≤ 0.05, ** p ≤ 0.01, *** p ≤ 0.001, **** p ≤ 0.0001. **(C)** Cross-sections of stage 10A follicles expressing E-Cad::GFP (green) treated with DMSO (control) or LatA. Single channels show E-Cad or Dextran (grey). Note enlarged intercellular gaps (magenta) on apical side in LatA-treated follicles. **(D)** Apical sections of mbFCs expressing E-Cad::GFP in stage 8, stage 9, and stage 10A follicles treated with DMSO (control) or 2 µM LatA for 45 minutes before imaging. Single channels show E-Cad (heatmap) and dextran (grey). Note absence of E-Cad from vertices and enlarged intercellular gaps in LatA-treated follicles. **(E)** Patency index on basolateral and apical sides of follicles treated with DMSO (control) or LatA. Stages and number of follicles (n) analyzed are indicated. P-values (Welch’s two-sample t-test): * p ≤ 0.05, ** p ≤ 0.01, *** p ≤ 0.001, **** p ≤ 0.0001. **(F)** Apical sections of mbFCs carrying *ssh^1-11^* clones (marked by absence of RFP (yellow)), stained for E-Cad (magenta) and p-MyoII (cyan). Note accumulation of p-MyoII at vertices at stage 10A (arrowhead). **(G)** Quantification of Phalloidin signals (normalized) at apical vertices in *ssh^1-11^* and control cells at stage 8 and stage 10A. Number of follicles (n) analyzed is indicated. P-values (Welch’s two-sample t-test): * p ≤ 0.05, ** p ≤ 0.01, *** p ≤ 0.001, **** p ≤ 0.0001. **(H)** Basolateral section of follicle carrying *ssh^1-11^* clone marked by absence of RFP (magenta). Dextran (green) marks intercellular spaces. **(I)** Patency index in *ssh^1-11^* clones and control cells. Number of follicles (n) analyzed is indicated. P-values (Welch’s two-sample t-test): * p ≤ 0.05, ** p ≤ 0.01, *** p ≤ 0.001, **** p ≤ 0.0001. **(J)** Scheme of Cofilin regulation. Phosphorylation by LIMK1 inactivates, while dephosphorylation by Slingshot activates, Cofilińs F-actin depolymerizing activity. Scale bars: (A) 1 µm (xy section), 5 µm (xz section); (C,H) 10 µm; (D,F) 5 µm. See also Figures S3 and S4.

### Enabled/VASP is dispensable for vertex-associated F-actin assembly in FCs

A key F-actin regulator, Enabled / vasodilator-stimulated protein (Ena/VASP), promotes assembly of vertex-associated F-actin in mammalian cells ^52^ and in *Drosophila* ^53,54^. A GFP::Ena fusion protein localized, besides cytoplasmic accumulations, to puncta abutting FC vertices in pre-patent follicles (Fig. S4A,B), resembling Jub, Vinc, and Zyx localization (Fig.1E, Fig. S1C,D), and consistent with direct binding of Ena to Vinc and Zyx ^55^. The GFP::Ena puncta disappeared from vertices by stage 9 (Fig. S4A,B), coinciding with a reduction of vertex-associated F-actin (Fig. S1A,B). To probe Enás role in organizing this F-actin pool, we depleted Ena function either by sequestrating Ena protein at mitochondria (Fig. S4D; ^53,56^) or by inducing FC clones homozygous for a newly generated *ena* deletion allele (*ena^1.8^*; Fig. S4C,E). Both led to depletion of Ena protein but had no or only slight effects on F-actin accumulation at apical vertices (Fig. S4D,E,G), and patency was not detectably affected (Fig. S4H,I). By contrast, Ena overexpression caused excessive F-actin accumulation and formation of actin-rich-protrusions at apical cell boundaries and vertices (Fig. S4F,G). Thus, although Ena can induce vertex-associated F-actin assembly in FCs, it is dispensable for this function, presumably due to redundancy with other factors.

### Hypertonic shock induces opening of vertices *ex vivo*

Searching for ways to analyze the dynamics of patency, we noticed that treatment of cultured stage 9 follicles with hypertonic medium induced transient opening of FC vertices, resembling the intercellular gaps observed during patency (Fig. 5A; ^29^). Addition of PBS or sucrose to the medium induced rapid formation of intercellular gaps, which reached maximal size after approximately one minute and closed after 5 minutes (n=5; Fig. 5A,B; Video S2). Addition of DMSO had a more gradual effect, and caused an initial contraction of cells, followed by opening of vertices with a peak in gap size at seven minutes and closure of gaps after 25 minutes (n=8; Fig. 5A,B; Video S2). By contrast, hypotonic treatment by diluting (1:1) the medium with water did not alter cell contacts at stage 9 (Fig. 5A,B; n=8) but induced closure of intercellular gaps in patent (stage 10A) follicles (Fig 5C,D; n=5). Interestingly, while opening of vertices upon DMSO-induced hypertonic treatment was consistently observed in vitellogenic (stage 9) follicles (Fig. 5A,B), only smaller and more variable openings were observed at earlier stages (Fig. 5E,F). In previtellogenic follicles (stage 5) with proliferating FCs, larger transient gaps formed at boundaries and cleavage furrows of dividing cells (Fig. 5E; Video S3), consistent with disengagement and remodeling of adhesion during mitosis ^57,58^. These findings suggest that hypertonic treatment can induce opening of FC vertices when remodeling of cell-cell junctions and of actomyosin has provided a permissive condition for initiation of patency.

**Figure 5.**
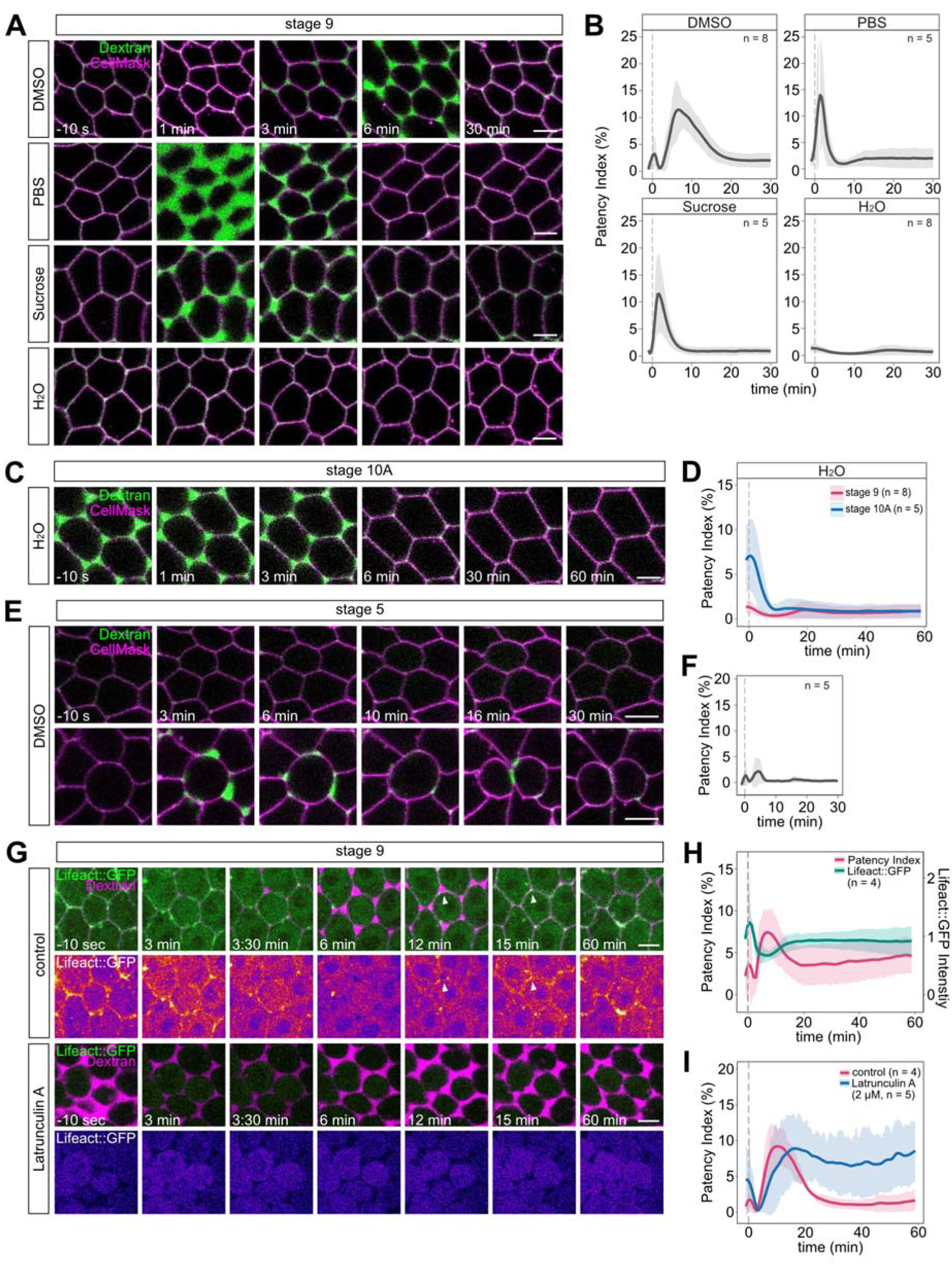
Hypertonic shock induces opening of follicle cell vertices *ex vivo*. **(A)** Stills (basolateral sections) from time-lapse movies of early patent (stage 9) follicles exposed to hypertonic (305 mosmol/L / 2.38% DMSO, 88,96 mosmol/L / 0.238x PBS, or 50 mosmol/L / 50 mM Sucrose) or hypotonic (50% H_2_O) medium at t=0 min. Dextran (green) marks intercellular spaces, CellMask (magenta) labels plasma membranes. **(B)** Patency index over time after hypertonic or hypotonic treatment as indicated. Graphs show mean values and standard deviations, smoothed using a running median. **(C)** Basolateral sections of patent (stage 10A) follicle exposed to hypotonic (50% H_2_O) medium at t=0 min. Note that intercellular gaps close after addition of H_2_O. **(D)** Patency index over time after hypotonic treatment of stage 9 (purple line) or stage 10A (blue line) follicles. **(E)** Hypertonic (DMSO) treatment of stage 5 follicles. Note that vertices of interphase cells remain closed after adding DMSO (top), whereas transient leaks appear at boundaries of dividing cells (bottom). **(F)** Patency index over time of stage 5 follicles. **(G)** Stage 9 follicles expressing Lifeact::GFP (driven by *CY2*-Gal4; green), treated with DMSO (control; top) or 2 µM LatA (bottom) for 45 minutes before imaging. Dextran (magenta) marks intercellular spaces. At t=0 min, follicles were exposed to DMSO-induced hypertonic shock. Note that Lifeact::GFP disappears before gap opening and relocalizes at vertices (arrowheads) when gaps start to close. Open vertices fail to close in LatA-treated follicles. **(H)** Quantification of Lifeact::GFP signals at cell boundaries (green line) and of patency index (purple line) over time in control follicles. **(I)** Patency index over time of control (purple line) and LatA-treated (blue line) follicles. Scale bars: (A, C, E, G): 5 µm. See also Figure S5 and Video S2, S3, S9 and S10.

Transient intercellular gaps after DMSO treatment were detectable along basolateral membranes (Fig. 5; Video S2) and at the level of AJs (Fig. S5A; Video S4). Interestingly, within 4 min after DMSO-induced hypertonic shock, MyoII::GFP redistributed from the cortex to the medial region as vertices opened (Fig. S5A-C; Video S5), resembling the behavior of MyoII in untreated follicles entering patency at stage 9 (Fig. 1D). Similarly, mNG::Rok transiently receded from vertices after DMSO treatment, coinciding with gap opening, and reappeared when vertices closed (Fig. S5D; Video S6). Consistent with these findings, intercellular gaps after DMSO treatment were enlarged and failed to close in Rok-depleted tissue (Fig. S5E; Video S7). Conversely, hyperactivation of MyoII in *flw^FP41^* clones prevented the opening of gaps after DMSO treatment (Fig. S5G; Video S8).

Given these similarities between patency and the response to DMSO-induced hypertonic shock, we employed this assay to probe the roles of F-actin in vertex opening and closure. Cortical F-actin (visualized by Lifeact::GFP) increased 10 sec after adding DMSO, coinciding with initial contraction of FCs (Fig. 5G; Video S9). Subsequently, F-actin disappeared from the cortex as intercellular gaps became visible and re-appeared at vertices when gaps started to close. Interestingly, Lifeact::GFP signals were anti-correlated with intercellular gap size: minimal signals concurred with maximal gap size and vice versa (Fig. 5H), suggesting that disassembly of cortical F-actin may be required for vertex opening, while F-actin assembly may promote closure of gaps.

### Cofilin-dependent F-actin disassembly is required for opening intercellular gaps

To test this hypothesis, we manipulated F-actin dynamics using pharmacological inhibitors or genetics and analyzed gap opening and closure following DMSO-induced hypertonic shock. In follicles treated with LatA, Lifeact::GFP signals were uniformly low and did not change after adding DMSO, although FCs contracted after 3 minutes (Fig. 5G,I; Video S10). Intercellular spaces in LatA-treated follicles opened, with gaps reaching maximal size after 16 minutes, but, unlike controls, failed to close within one hour (Fig. 5G,I), consistent with a requirement of F-actin assembly for vertex closure.

Conversely, FC clones lacking the protein phosphatase Slingshot (Ssh; ^37^), which promotes F-actin disassembly by dephosphorylating and thereby activating Cofilin, showed substantial accumulation of apical F-actin that persisted into stage 10A (Fig. 4F,G). Interestingly, *ssh^1-11^* mutant FC clones displayed reduced intercellular openings during patency (Fig. 4H,I). These results indicate that Slingshot/Cofilin-dependent F-actin disassembly is required for vertex opening, while F-actin assembly promotes closure of gaps after patency.

### Acute activation of Rho signaling is sufficient to prevent patency

Our findings suggest that Rho-induced actomyosin contractility is necessary to maintain FC vertices closed. To test whether Rho signaling is also sufficient to prevent vertex opening, we combined the hypertonic shock assay (Fig. 5) with the CRY2/CIBN optogenetic system ^59^ to acutely activate Rho signaling (Fig. 6A). We generated mosaic follicles co-expressing CRY2-RhoGEF2::mCherry and plasma membrane-anchored CIBN::pmGFP in FC clones (Fig. 6B). Illumination with blue light (488 nm) triggered rapid plasma membrane recruitment of CRY2-RhoGEF2::mCherry. When a hypertonic shock was applied by adding DMSO nine minutes after optogenetic Rho activation, control cells opened their vertices normally, whereas vertex opening was strongly reduced in FCs with activated Rho signaling (Fig. 6B,C; Video S11). As a control, chronic membrane localization of Rho in flies expressing CRY2-RhoGEF2::mCherry and CIBN::pmGFP under constant light blocked patency, whereas CRY2-RhoGEF2::mCherry expressed alone was distributed in the cytoplasm and did not affect patency (Fig. 6D,E). Thus, the blockade of patency after optogenetic Rho activation is not caused by chronic RhoGEF2 overexpression, and acute activation of Rho-induced actomyosin contractility is sufficient to prevent opening of FC vertices.

**Figure 6.**
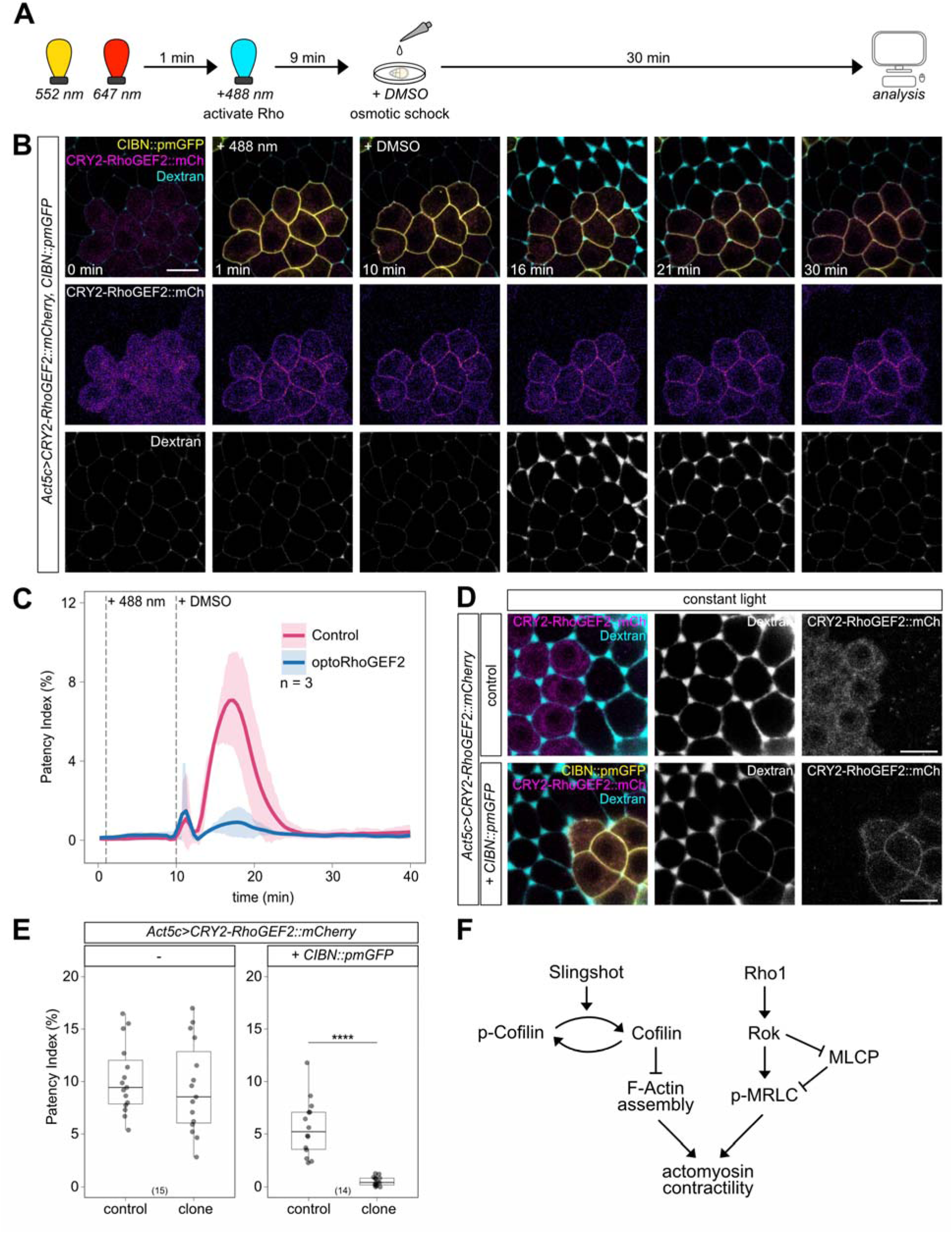
Acute activation of Rho is sufficient to prevent patency. **(A)** Scheme of combined Rho activation and hypertonic shock experiment. Follicles are exposed to blue light (488 nm, 9 min) to activate Rho signaling before adding DMSO to induce hypertonic shock at t=10 min. **(B)** Stills (basolateral sections) from time-lapse movie of stage 9 follicle expressing CRY2-RhoGEF2::mCherry (magenta) and CIBN::pmGFP (yellow) in FC clone. Dextran (cyan) marks intercellular spaces. Blue light at t=1 min (+488 nm) triggers membrane localization of CRY2-RhoGEF2::mCherry. DMSO-induced hypertonic shock at t=10 min triggers vertex opening in control tissue, whereas vertices of optoRhoGEF-expressing cells do not open. Single channels show CRY2-RhoGEF2::mCherry intensities (heatmap) and dextran (grey). **(C)** Patency index over time of control cells (purple line) and optoRhoGEF-expressing cells (blue line). Mean values and standard deviations (smoothed using a running median) are shown. **(D)** Basolateral sections of stage 10A follicles from females raised in constant light, expressing CRY2-RhoGEF2::mCherry (magenta) either alone (top) or with CIBN::pmGFP (yellow, bottom) in FC clones. Note that continuous Rho activation by plasma-membrane-targeted RhoGEF2, but not overexpressed RhoGEF2 alone, blocks vertex opening. **(E)** Patency index of FCs expressing CRY2-RhoGEF2::mCherry either alone (left) or together with CIBN::pmGFP (right) under constant light. Act5c>CRY2-RhoGEF2::mCherry-expressing cells (“clone”) are compared to surrounding cells (“control”). Number of follicles (n) analyzed is indicated. P-values (Welch’s two-sample t-test): * p ≤ 0.05, ** p ≤ 0.01, *** p ≤ 0.001, **** p ≤ 0.0001. **(F)** Regulation of actomyosin contractility in FCs at patency initiation. Scale bars: (B, D) 10 µm. See also Video S11.

## Discussion

We used follicular patency in *Drosophila* as a model to understand how an intact epithelial barrier is transiently breached by opening of TCJs to enhance paracellular permeability. Our work reveals that this process is governed by modulation of actomyosin contractility at cell vertices. We show first that before patency, apical vertex-anchored actomyosin bundles generate tensile forces that keep vertices closed. Sustained MyoII activity in pre-patent follicles depends on Rho-associated kinase, which accumulates at vertices along with tension-sensing junctional proteins Jub, Vinc, and Zyx. Second, when TCJ opening initiates, MyoII redistributes from vertex-anchored peripheral bundles to a medial pool, Rok and tension-sensing proteins withdraw from vertices, and junctional tension is released. Third, this transition requires inactivation of MyoII by Myosin light-chain phosphatase (MLCP), which is normally inactivated through phosphorylation by Rok. Removal of Rok at the onset of patency abrogates inactivation of MLCP, enabling it to inactivate MyoII, thereby releasing constricting forces and allowing vertices to open. Finally, we show that a dynamic (LatA-sensitive) pool of F-actin is associated with apical FC vertices and helps to keep vertices closed. Initiation of patency requires Slingshot/Cofilin-dependent disassembly of this F-actin network. Conversely, F-actin assembly is required for closing intercellular gaps after patency.

Thus, the opening of intercellular gaps in the follicular epithelium does not require actomyosin-based pulling forces but depends on relaxation of actomyosin contractility, which acts to constrict vertices. This is consistent with a force transduction model in which TCJ integrity is maintained by contractile actomyosin bundles that are anchored in an end-on fashion at vertices (Fig. 1G; ^19,42–44^). This type of actomyosin arrangement was described in mammalian mammary epithelial EpH4 cells, where the tricellular tight junction (tTJ) protein Tricellulin regulates F-actin organization through Cdc42 ^60^ and anchors actin filaments at vertices via α-Catenin, which in turn recruits Vinculin to reinforce linkage to the tTJ-associated actin network ^52^. Resembling our findings in *Drosophila*, MyoII is concentrated at tTJs and is required for maintaining tTJ integrity in EpH4 cells ^52^. While Tricellulin is not conserved in *Drosophila*, the vertex-specific adhesion protein Sidekick (Sdk) plays an analogous role by recruiting MyoII ^15^, suggesting that MyoII has a common function in TCJ maintenance in vertebrate and invertebrate epithelia.

### Tension-sensitive actomyosin assembly at cell vertices

What mediates the assembly of vertex-anchored F-actin? Tension-sensitive Vinculin recruitment promotes F-actin organization at vertices and is required for TCJ integrity and barrier function in mammalian cells ^52^ and in the *Xenopus* epidermis ^61^. Although Vinculin also accumulates at vertices of pre-patent follicles, we found Vinculin to be dispensable for F-actin organization and for control of patency (not shown), consistent with earlier reports of *vinc* being a non-essential gene in *Drosophila* ^62,63^. This could be explained by functional redundancy of Vinculin with other proteins that mediate tension-sensitive linkage between AJs and F-actin, such as Ajuba and Zyxin ^64^. Interestingly, Vinculin and Zyxin recruit the actin polymerase Ena/VASP ^55^ to AJs under tension, where Ena promotes F-actin assembly ^65^. Ena is recruited to vertices in *Drosophila* (Fig. S4; ^53,54,66^) and in mammalian epithelial cells^52^. However, like Vinculin, Ena was not essential for F-actin assembly in FCs, suggesting that Ena acts in a redundant fashion along with other factors that ensure robust maintenance of this F-actin network.

### Dual regulation of actomyosin contractility during patency

Work in mammalian cells suggested that tensile forces sustain a positive feedback loop that generates vertex-anchored actin filaments through tension-dependent recruitment of Vinculin and actin-polymerizing proteins, such as Ena/VASP ^52,65^. We showed that local activation of MyoII by vertex-localized Rok generates contractile forces, which in turn may promote tension-dependent F-actin assembly at FC vertices. Our genetic results indicate that this feedback loop can be broken by inactivating MyoII through MLCP or by promoting Ssh/Cofilin-dependent F-actin disassembly (Fig. 6F). Thus, patency initiation involves dual regulation of actomyosin contractility, and both, MyoII inactivation and F-actin disassembly, are required for normal vertex opening. Interestingly, we identified two phosphatases, MLCP and Slingshot, as essential regulators of these processes during patency initiation. It will be interesting to elucidate whether Rho/Rok signaling regulates both phosphatases to jointly control F-actin dynamics and Myosin activity, or whether different upstream signals are involved. While Rok clearly controls MyoII activity via MLCP in pre-patent FCs, our genetic results suggest that the conserved parallel pathway downstream of Rok that regulates F-actin dynamics via LIMK1-dependent inactivation of Cofilin ^34,36,67^ is not active in FCs. Instead, we found that the phosphatase Ssh is essential for Cofilin-dependent F-actin disassembly at the onset of patency. As Ssh has not been reported to be regulated by Rho signaling, it will be interesting to elucidate how Ssh activity or localization is controlled in FCs. Interestingly, the mammalian Ssh homolog SSH1L can antagonize Rok signaling by dephosphorylating the Rok substrate MyoII phosphatase MYPT-1 ^68^. While it is not known whether Ssh acts analogously in *Drosophila*, such a mechanism could mediate coordinate regulation of actin disassembly and Myosin inactivation in patency initiation.

Although the mechanisms underlying gap closure and restoration of barrier function after patency remain to be characterized in detail, our findings suggest that F-actin reassembly aids in sealing intercellular gaps to terminate patency. Similarly, localized transient Rho activation mediates actin polymerization and Rok-dependent contractility to repair tight junction leaks in *Xenopus* epidermis ^7^. How Rho activity is spatially and temporally regulated to restore barrier integrity, or to control TCJ opening, as in the case of patency, is not clear. In vertebrate endothelial cells, inflammatory mediators, such as histamine, thrombin, or VEGF, induce opening of cell-cell junctions to increase vascular permeability by activating actomyosin contractility through RhoA ^69^. Thus, in this system, unlike *Drosophila* follicles or *Xenopus* epidermis, Rho-induced actomyosin contractility increases junctional permeability rather than preventing it. In endothelial cells, which differ from cuboidal epithelia in cell shape and cytoskeletal organization, Rho activation induces radially aligned stress fibers that exert pulling and shear forces on VE-Cadherin-based junctions, resulting in junctional leaks ^70,71^. By contrast, contractility of vertex-anchored actomyosin filaments in FCs maintains TCJ integrity and helps to prevent premature TCJ opening. Thus, the cell-type-specific organization of actomyosin at cell vertices determines the mode of contractility-dependent regulation of epithelial permeability.

## RESOURCE AVAILABILITY

### Lead Contact

Further information and requests for resources and reagents should be directed to the Lead Contact, Stefan Luschnig.

### Materials Availability

*Drosophila* strains described in this study are available from the Bloomington *Drosophila* stock Center (BDSC), the Kyoto *Drosophila* Genetic and Genomic Resources Center (DGGR), or from the Lead Contact. Antibodies are available from the sources listed in the Key Resources Table.

### Data and Code Availability

All data reported in this paper will be shared by the lead contact upon request. This paper does not report original code.

Any additional information required to reanalyze the data reported in this paper is available from the lead contact upon request.

## Acknowledgements

We thank Wilko Backer for expert technical help, Konrad Basler, Yohanns Bellaïche, Nick Brown, Stefano De Renzis, Katja Röper, Antoine Guichet, Veit Riechmann and Frank Schnorrer for providing antibodies and fly stocks, and Kiryl Piatkevich for sharing the mBaoJin plasmid. We thank Archana Rajan Vellandath for help with osmotic shock experiments. We acknowledge service by the Münster Imaging Network, the Developmental Studies Hybridoma Bank (created by the NICHD of the NIH and maintained at the University of Iowa), and the Bloomington Drosophila Stock Center (NIH P40OD018537). This work was supported by the Deutsche Forschungsgemeinschaft (SFB 1348 “Dynamic Cellular Interfaces”), the “Cells-in-Motion” Cluster of Excellence (EXC 1003-CiM), and the University of Münster.

## Author contributions

Conceptualization, T.J., J.I.S., S.L.; Methodology, all authors; Investigation, all authors; Formal Analysis, T.J., J.I.S., S.R.; Visualization, T.J., J.I.S., S.R.; Reagents and tools, all authors; Figures, T.J., S.L.; Writing – Original Draft, T.J.; Writing – Review and Editing, S.L.; Funding Acquisition, S.L; Supervision, S.L.

## Materials and Methods

### EXPERIMENTAL MODEL AND SUBJECT DETAILS

#### *Drosophila* strains and genetics

Flies were reared at 22°C on standard cornmeal-molasses-yeast food. Unless noted otherwise, *Drosophila* strains are described in FlyBase. Genotypes of specimens shown in the figures are listed in Table S1. All strains used in this work are listed in the Key Resources Table.

Mosaic follicles carrying *ena*^1^*^.8^*, *flw ^FP41^*, or *ssh^1-11^* mutant clones were generated using Flipase-induced mitotic recombination. A strain carrying a *hs-Flp* source and the appropriate FRT chromosome marked with either *ubi-GFP-nls* or *ubi-mRFP-nls* transgenes was crossed to a strain carrying the mutation of interest on the corresponding FRT chromosome. To induce Flp-mediated mitotic recombination, progeny of these crosses was heat-shocked at the pupal stage (48-96 hours after puparium formation) for 3 x 60 mins at 37°C in a water bath. For RNAi-mediated knockdown or overexpression experiments, flip-out clones were generated by crossing flies carrying *hs-Flp^122^* and the UAS construct of choice to flies carrying an *act5C>CD2>Gal4* flip-out construct. Progeny of these crosses was heat shocked as adults (1-2 days after eclosion) for 30 min at 37°C in a water bath.

#### Generation of new *ena* alleles

A transgenic CRISPR approach was used to generate *ena* deletion alleles. Male flies expressing from a transgene (B-83021) two different guide RNAs (sgRNA-WKO.1-C12.1, sgRNA-WKO.1-C12.2; ^86^) targeting the *ena* coding sequence were crossed to females expressing *nos*-Cas9 in the germline. F1 males expressing *nos-Cas9* and the two guide RNAs were crossed with *y w; Gla/CyO Dfd-GMR-YFP* females to establish stocks. Single F2 males were mated with *ena*^23^*/CyO* (B-25405) females to test for complementation of *ena*^23^ lethality. Lines were established from crosses that failed to complement lethality. The *ena^1.8^* mutation was recombined onto an FRTG13 chromosome for subsequent mosaic analyses. To identify lesions in the *ena* locus of newly induced *ena* alleles, genomic DNA was isolated from homozygous embryos (detected by the absence of YFP signal from the *CyO Dfd-GMR-YFP* balancer chromosome) and the *ena* genomic region was amplified using oligonucleotides (5’-CCTGGTACACATTCGCGTTT and 5’-TGTGCAGCTGAGATCTTGGA) and sequenced.

#### Transgenic constructs

The UAS-Lifeact::mBaoJin construct was generated as follows. The Lifeact peptide ^92^ coding sequence was produced by annealing two complementary oligonucleotides (5’-CAAAATGGGCGTGGCCGATCTGATCAAGAAGTTCGAGAGCATCAGCAAGGAGGAA and 5’-CCGGTTCCTCCTTGCTGATGCTCTCGAACTTCTTGATCAGATCGGCCACGCCCATTTTG GTAC), resulting in a dsDNA fragment with overhangs compatible with AgeI and XbaI restriction sites, respectively. A plasmid carrying the mBaoJin ^89^ coding sequence was a gift from Kiryl Piatkevich (Westlake University, Zhejiang, China). The mBaoJin coding sequence was amplified using forward (5’-GAACCGGTCGCCACCGTGTCGAAAGGTGAAGAGGA) and reverse (5’-GATCTAGATTATTTGTACAGCTCGTCCATGC) primers and the product was cut with AgeI and XbaI. The Lifeact and mBaojin fragments were co-ligated with pUASt-attB ^88^ after cutting with AgeI and XbaI. The pUASt-Lifeact::mBaoJin construct was integrated into the attP40 and attP2 landing sites, respectively, using PhiC31-mediated site-specific integration ^88^. All plasmids used in this study are listed in the Key Resources Table.

#### Antibodies and immunostainings

Female flies were collected 24-48 hours after eclosion and were fed on bakeŕs yeast for two days before dissecting ovaries in Shields and Sang M3 medium (Biomol; supplemented with 1x Penicillin/Streptomycin and 10% fetal bovine serum; M3++ medium). For immunostaining, dissected ovarioles were fixed in 4% PFA in PBS for 15 minutes at room temperature, rinsed quickly 3 times in PBT and blocked 3 x 30 minutes in 10% BSA in PBT (0.2% Tween-20). Primary antibodies (rat anti E-Cad, 1:50, DSHB; mouse anti Ena, 1:50, DSHB) were diluted in 10% BSA in PBT and incubated with slow agitation overnight at 4°C. After 3 washes in PBT, samples were blocked 3 x 30 mins in 10% BSA/PBT. Fluorophore-conjugated secondary antibodies were added at 1:500 in 10% BSA/PBT and incubated for 2.5 hours at room temperature or overnight at 4°C. After six washes (10 min each) in PBT, samples were mounted in ProLong Gold antifade medium (Thermo Fisher).

For anti-phospho-Myosin immunostainings, follicles were dissected in M3++ medium and then fixed in 4% PFA in PBS supplemented with phosphatase inhibitor solution (modified after ^93^). Subsequent steps were done as described above, except that primary antibodies (rabbit anti p-MyoII 1:50, Cell Signaling Technologies) were incubated for 24 hours at 4°C. Phosphatase inhibitor solution was prepared as a 20x stock solution (modified after ^93^: 20 mM NaF, 20 mM β-Glycerophosphate, 20 mM Na_3_VO_4_, 100 mM Na_4_P_2_O_7_ • 10H_2_O).

F-actin in fixed samples was visualized using Phalloidin AlexaFluor 488 or Phalloidin AlexaFluor 568 (1:1000; Fisher Scientific), together with secondary antibodies.

#### Culture and imaging of ovarian follicles

For live imaging, follicles were dissected and cultured in M3++ medium. For plasma membrane staining, CellMask Orange Plasma Membrane Stain (6.25 µg/mL final concentration; Thermo Fisher) was added to the dissection medium. For permeability assays, follicles were transferred to 200 µL M3++ medium containing dextran (10 kDa) conjugated with AlexaFluor 647 (37.5 µg/mL, Life Technologies). For visualizing F-actin with Lifeact-Halo2, follicles expressing *UAS-Lifeact-Halo2* ^84^ driven by *GR1*-Gal4 were incubated in M3++ medium containing 1 µM HALO-Tag Ligand-Janelia Fluor-646 (Promega, GA1120) for 20 minutes before mounting. Follicles were imaged immediately after mounting on glass slides with cover slips.

For osmotic treatment, dissected follicles were transferred to 8-well glass-bottom chambers (VWR 734-2061) coated with Poly-D-Lysine (1 mg/mL in 0.15 M H_3_BO_3_ pH 8.4). Each chamber contained 200 µL M3++ medium supplemented with dextran-AlexaFluor 647 as above. Osmolarity was changed during imaging by adding the respective osmotic agent as follows: 10 µL 50% DMSO in M3++ (final concentration 2.38% DMSO or 304 mosmol/L); 10 µL 5x PBS in M3++ (final concentration 0.238x PBS or 88,96 mosmol/L); 50 µL 250 mM Sucrose in M3++ (final concentration 50 mM or 50 mosmol/L). For hypotonic shocks, 150 µL H_2_O were added to follicles in 150 µL medium containing dextran-AlexaFluor 647. The concentration of dextran-AlexaFluor 647 (37.5 µg/mL) was kept constant during osmotic shocks by adding the appropriate amount of dextran-AlexaFluor 647 along with the osmotic reagent.

For Latrunculin A treatment, follicles were dissected and incubated for 45 minutes in M3++ medium supplemented with Insulin (0.4 mg/mL; Sigma I0516) and Latrunculin A (2 µM; Abcam #ab144290).

For long-term high-resolution live imaging, follicles were mounted in fibrinogen gel as described in ^94^. Follicles were dissected in serum-free M3 medium containing Fibrinogen (5 mg/mL; Sigma F3879) and transferred in 7 µL of medium to a MatTek round dish imaging chamber (MatTek Corp., P35G-1.5-14-C). Fibrinogen was clotted by adding 1.6 µL Thrombin (1000 units/mL; Sigma T4648) for 5 to 10 minutes. The clot was covered with 200 µL Schneider’s medium supplemented with 20% fetal bovine serum, Glucose (1 g/L) and Insulin (0.4 mg/mL; Sigma I0516; adapted from ^94^).

For osmotic shock experiments, follicles were transferred to 8-well glass-bottom chambers (VWR 734-2061) coated with Poly-D-Lysine (1 mg/mL in 0.15 M H_3_BO_3_ pH 8.4). Each chamber contained 200 µL M3++ medium supplemented with 10 kDa dextran-AlexaFluor647 as above.

For optogenetic experiments, flies were fed on yeast in constant darkness at 25°C for two days before dissection. Flies were dissected and samples were mounted under red light before imaging on a Leica SP8 confocal microscope.

For all remaining experiments, follicles were mounted in M3++ medium (containing dextran-AlexaFluor647 where applicable) on glass slides with cover slips held by spacers (a cut cover slip between two layers of double-sided tape).

All media and reagents are listed in the Key Resources Table.

#### Laser Ablations

Dissected follicles were mounted in eight-well glass-bottom chambers coated with poly-D-Lysine and covered with 200 µL M3++ medium. Imaging was performed on a Zeiss Spinning Disk (Yokogawa CSU-X1) confocal microscope with 100x / NA 1.45 oil immersion objective. Images of a single z-plane were acquired at a rate of two frames per second over 40 seconds. Ablations were performed using a pulsed 355 nm laser (Rapp OptoElectronics). Vertices were ablated within a 5-pixel ROI with one laser pulse (530 ms) per vertex. Vertex displacement was tracked in FIJI using the Manual Tracking Plugin (https://imagej.net/ij/plugins/track/track.html). Using R (v4.2.3, with RStudio Interface (2023.03.0+386)), the displacement of the three vertices next to the ablated vertex was calculated as Euclidean distance 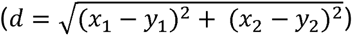. The mean displacement and standard deviation of the three vertices per movie were calculated and the time series data was smoothed by applying a running median with a median window of k=5.

#### Microscopy

Confocal images were acquired on a Leica SP8 confocal microscope with 40x / NA 1.3 or 63x / NA 1.4 oil immersion objectives or on a Zeiss LSM980 confocal microscope equipped with an Airyscan 2 module and a 63x / NA 1.4 oil immersion objective. Airyscan images were processed using Zeiss Zen blue (v3.5.093.00010) Software with super-resolution filter set to 6.3. Image data acquired with the resonant scanner (Leica SP8) was deconvolved using Leica Lightning Deconvolution (v3.5.7.23225).

### QUANTIFICATION AND STATISTICAL ANALYSIS

#### Image analysis

Images were processed using FIJI/ImageJ (v1.42; ^90^) and OMERO (5.8.0). Image panels were assembled using OMERO.figure v6.2.2 ^95^ and Affinity Designer (v2.3.0).

To measure patency index, confocal sections of main body FCs were acquired at 25% of total epithelium height from the basal surface. A median filter was applied to the dextran-AlexaFluor647 channel, and a threshold was set to create a mask of the dextran signal. The Analyze Particle feature in FIJI was used to measure the area filled with dextran signal. The patency index represents the total area of dextran-filled spaces within the ROI divided by the total ROI area ^29^.

For quantification of signal intensities at follicle cell vertices, average projections of 20 confocal sections spanning 3 µm of the apical region were generated. Fluorescence intensities at vertices were measured within circular ROIs with 60 pixels diameter (63 x objective) or 55 pixels diameter (40 x objective) centered on a vertex. Background was subtracted and mean intensities were normalized to the mean intensities at the earliest stage analyzed.

For quantification of Lifeact::GFP signals at the cortex in time-lapse movies, the plasma membrane of the most apical section at every time point was segmented using the CellMask channel to create a mask. The mask was dilated twice (using the Dilate function in Fiji) and reapplied to the Lifeact::GFP channel to measure intensities at every time point. The area filled with dextran signal was determined on the same confocal section.

For quantification of the ratio of MyoII::GFP signals between the cortex and the medial region in osmotic shock time-lapse movies, an average intensity projection of 14 slices (6 µm) was generated. The E-Cad::RFP channel was used to identify cortex and medial regions by segmenting bleach-corrected E-Cad::RFP signals at every time point. The medial region is the inverse and dilated mask of the cortical region. To measure dextran area in these movies, the most basolateral slice of each movie was analyzed as described above.

#### Statistics

Statistical analyses were performed in R (4.2.3) using RStudio Interface (2023.03.0+386). For phenotypic analyses, sample size (n) was not predetermined using statistical methods, but was chosen by considering the variability of a given phenotype, determined by the standard deviation. Sample size for each experiment is indicated in the graphs or figure legends. Experiments were considered independent if specimens were derived from different parental crosses. Investigators were not blinded to allocation during experiments. The data was tested for normality using the Shapiro-Wilk test. Welch’s two-sample t-test was used for normally distributed data. For data that was not normally distributed, or data that was compared between multiple stages, the Wilcoxon rank-sum test (R standard package) was used. P-values were corrected for multiple testing using the Bonferroni-Holm method ^96^.

## Supplemental information

Document S1. Figures S1–S5 and Table S1.

Video S1. Laser ablations of apical FC vertices. Related to Figure 1 and S2.

Video S2. Osmotic treatment of cultured follicles. Related to Figure 4 and 5.

Video S3. Hypertonic treatment of stage 5 follicles containing cells in interphase and mitotic cells. Related to Figure 4 and 5.

Video S4: Hypertonic treatment of follicle expressing E-Cad::RFP. Related to Figure 5.

Video S5: Hypertonic treatment of follicle expressing MyoII::GFP and E-Cad::RFP. Related to Figure 5.

Video S6: Hypertonic treatment of follicle expressing mNG::Rok. Related to Figure 5.

Video S7: Hypertonic treatment of mosaic follicle carrying Rok-depleted clone. Related to Figure 5.

Video S8: Hypertonic treatment of mosaic follicle carrying *PP1β* mutant clone. Related to Figure 5.

Video S9. Hypertonic treatment of follicle expressing Lifeact::GFP. Related to Figure 4 and 5.

Video S10. Hypertonic treatment of follicle expressing Lifeact::GFP treated with Latrunculin A. Related to Figure 4 and 5.

Video S11. Optogenetic activation of Rho signaling followed by hypertonic treatment. Related to Figure 6.

