## Supplemental Information for "Rho/Rok-dependent regulation of actomyosin contractility at tricellular junctions restricts epithelial permeability in Drosophila"

**Figure S1**

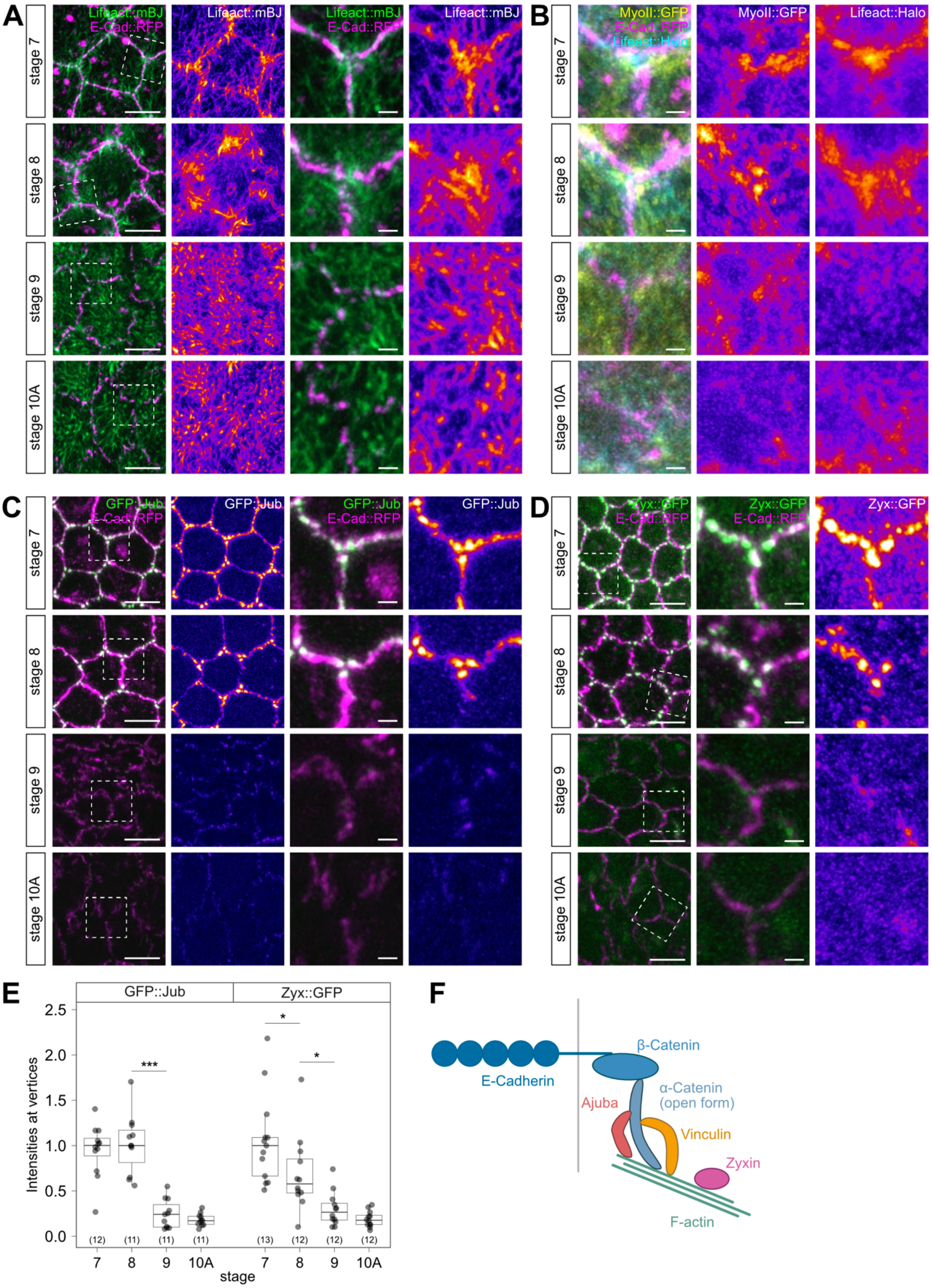
A
Lifeact::mBJ
E-Cad::RFP
stage 7
stage 8
stage 9
stage 10A
B
MyoII::GFP
Lifeact::Halo
C
GFP::Jub
D
Zyx::GFP
E
Intensities at vertices
stage
***
*
F
E-Cadherin
β-Catenin
α-Catenin (open form)
Ajuba
Vinculin
Zyxin
F-actin

**Figure S1 (related to Figure 1 and Figure 2)**

**Distribution of mechanosensitive proteins at follicle cell vertices.**

**(A)** Apical sections of mbFCs expressing endogenous E-Cad::RFP (magenta) and Lifeact::mBaoJin (driven by *tj*-Gal4; green). Single channels show Lifeact::mBaoJin intensities color-coded using heat map. Intensities are not comparable between stages in this panel but were adjusted for best visualization. Close-up views in (A) show single vertices marked by rectangles in overview.

**(B)** Close-up of single vertices in apical sections of mbFCs expressing E-Cad::RFP (magenta), MyoII::GFP (yellow) and Lifeact::Halo (conjugated with HaloTag-647, driven by *GR1*-Gal4; cyan). Single channels show MyoII and Lifeact::Halo intensities color-coded using heat map. Note that intensities are not comparable between stages in this panel but were adjusted per stage for best visualization.

**(C)** Apical sections of mbFCs expressing E-Cad::RFP (magenta) and GFP::Jub (green). Close-up views show single vertices. Note that Jub (intensities color-coded using heatmap) accumulates near vertices at stage 7 and 8, but not in stage 9 and 10A.

**(D)** Apical sections of mbFCs expressing endogenous E-Cad::RFP (magenta) and Zyx::GFP (green). Note that Zyx (intensities color-coded using heatmap) accumulates near vertices at stage 7 and 8, is reduced at stage 9, and absent at stage 10A.

**(E)** Quantification of GFP::Jub and Zyx::GFP signals (normalized) at apical vertices. Number of follicles (n) analyzed is indicated. P-values (Pairwise Wilcoxon Rank Sum Test): * $p \leq 0.05$, ** $p \leq 0.01$, *** $p \leq 0.001$, **** $p \leq 0.0001$.

**(F)** Scheme of tension-sensitive proteins at AJs. Vinculin and Ajuba bind to α-Catenin in its open conformation and to F-actin, while Zyxin associates with F-actin.

Scale bars: (A) 5 µm, close-ups 1 µm, (B) 1 µm, (C-D) 5 µm, close-ups 1 µm

**Figure S2**

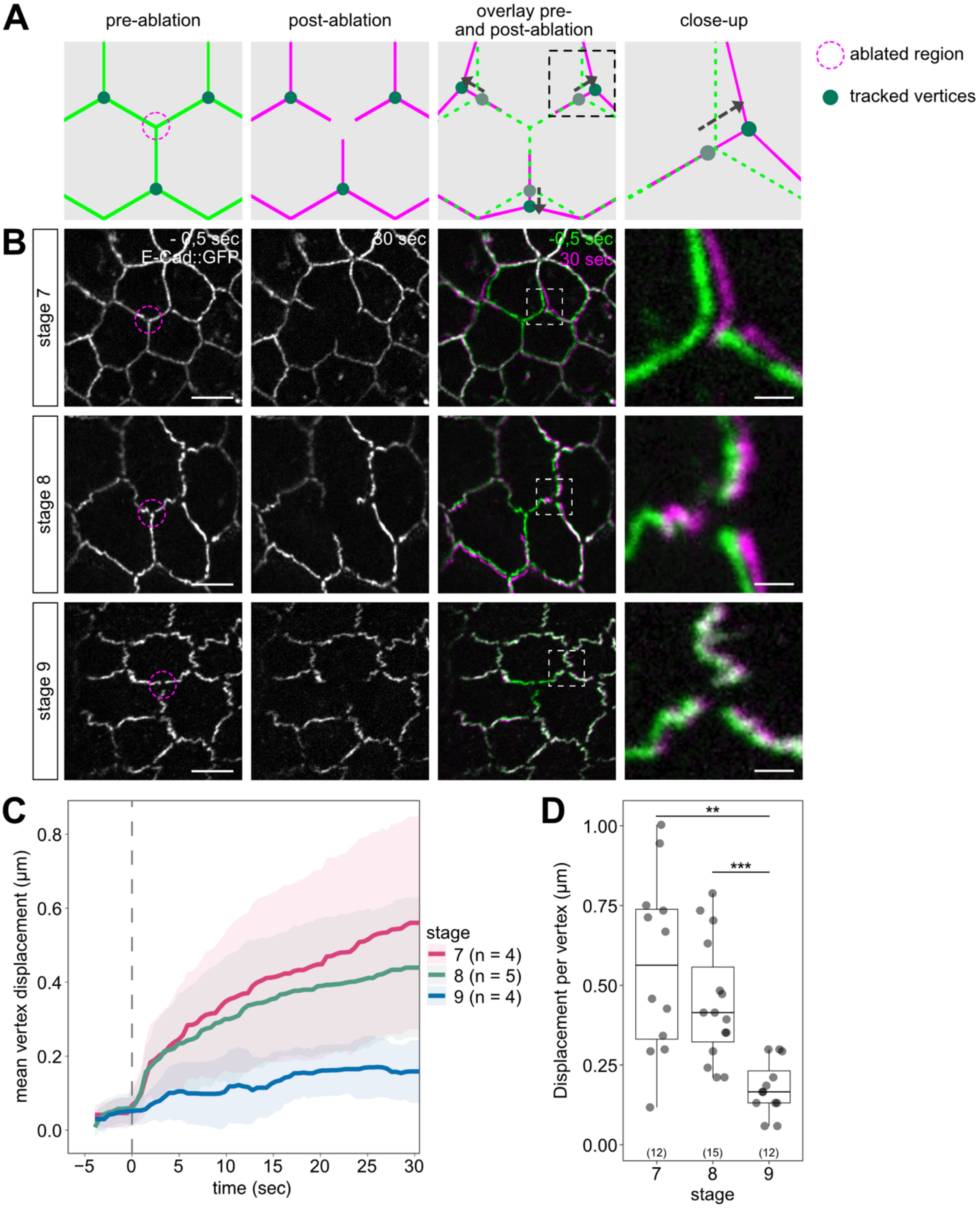
A
pre-ablation
post-ablation
overlay pre- and post-ablation
close-up
ablated region
tracked vertices
B
stage 7
stage 8
stage 9
- 0,5 sec
E-Cad::GFP
30 sec
-0,5 sec
30 sec
C
mean vertex displacement (µm)
time (sec)
stage
7 (n = 4)
8 (n = 5)
9 (n = 4)
D
Displacement per vertex (µm)
stage
**
***
(12)
(15)
(12)

**Figure S2 (related to Figure 1 and Figure 2)**

**Tension on vertices decreases at the onset of patency.**

**(A)** Schematic representation of UV laser ablation experiments and analysis. Green membranes represent last timepoint before ablation, magenta membranes represent timepoint 30 seconds post ablation.

**(B)** Apical vertex ablation in stage 7, 8 and 9 of follicles expressing E-Cad::GFP (grey), magenta ROI indicates ablated vertex. Merge panels show overlay of last frame before ablation (green) and frame 30 seconds after ablation (magenta). Close-up views show single vertices.

**(C)** Quantification of vertex displacement (normalized) upon ablation. Stage 7 (purple) show highest vertex displacement, while stage 9 (blue) shows lowest vertex displacement. Mean values and standard deviations smoothed using a running median are shown.

**(D)** Quantification of vertex displacement 30 seconds after ablation. Each datapoint represents one vertex. Number of follicles (n) analyzed is indicated. P-values (Pairwise Wilcoxon Rank Sum Test): * $p \leq 0.05$, ** $p \leq 0.01$, *** $p \leq 0.001$, **** $p \leq 0.0001$.

Scale bars: (B): 5 μm, close-ups 1 μm.

See also Video S1.

### Figure S3

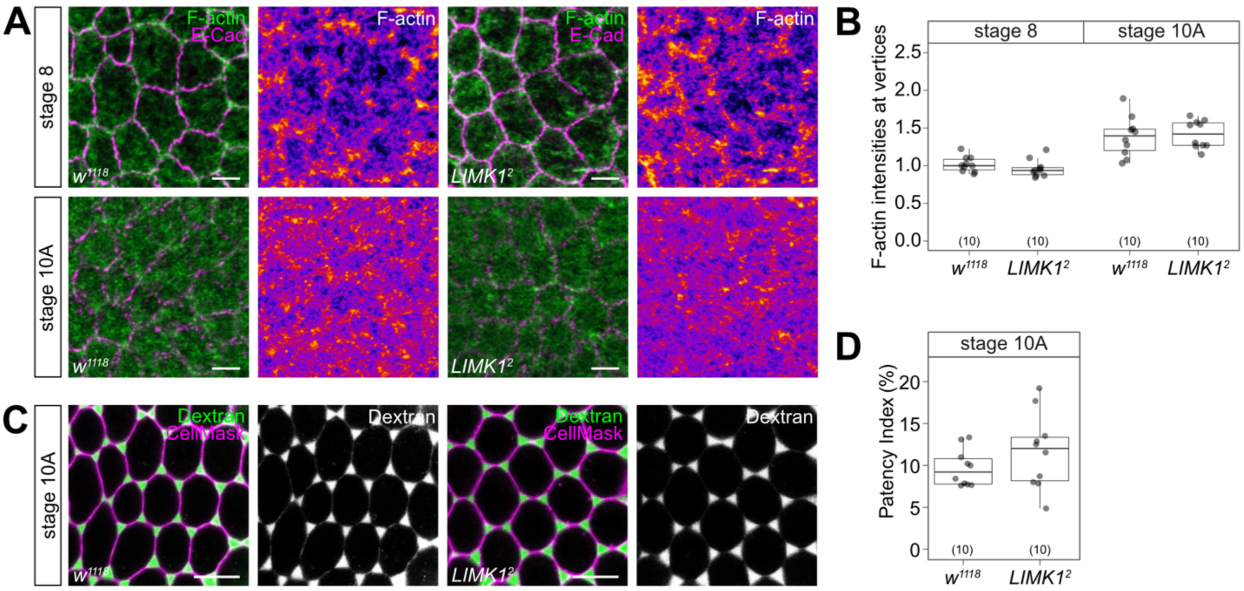


**Figure S3 (related to Figures 2, 3, and 4)**

**LIM domain kinase 1 is dispensable for F-actin remodeling and vertex opening in the follicle epithelium.**

**(A)** Apical sections of follicles from *w*$^{1118}$ (control, left) and *LIMK1*$^{2}$ homozygous females (right) stained for E-Cad (magenta) and F-actin (Phalloidin; green). Single channel shows Phalloidin intensities color-coded using heat map.

**(B)** Quantification of Phalloidin signals (normalized) at apical vertices in wild-type and *LIMK1*$^{2}$ follicles in stage 8 and stage 10A. Number of follicles (n) analyzed is indicated. P-values (Welch's two-sample t-test): * p ≤ 0.05, ** p ≤ 0.01, *** p ≤ 0.001, **** p ≤ 0.0001.

**(C)** Basolateral sections of *w*$^{1118}$ (control, left) and *LIMK1*$^{2}$ (right) follicles (stage 10A) incubated in fluorescent dextran (green). CellMask (magenta) labels plasma membranes.

**(D)** Quantification of patency index of mbFCs in wild-type and *LIMK1*$^{2}$ follicles (stage 10A). Number of follicles (n) analyzed is indicated. P-values (Welch's two-sample t-test): * p ≤ 0.05, ** p ≤ 0.01, *** p ≤ 0.001, **** p ≤ 0.0001.

Scale bars: (A) 5 µm, (B) 10 µm.

**Figure S4**

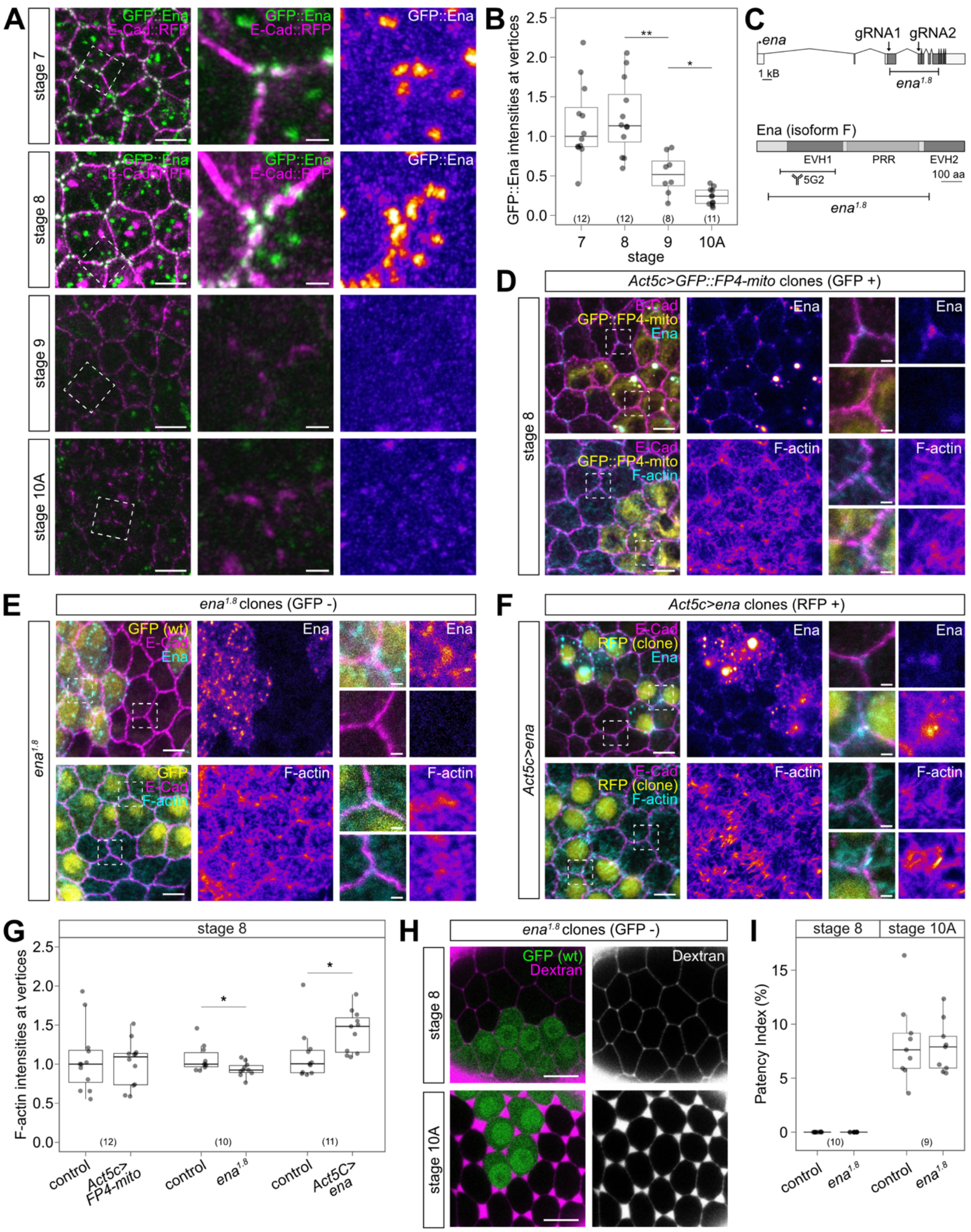
A
stage 7
stage 8
stage 9
stage 10A
GFP::Ena
E-Cad::RFP
B
GFP::Ena intensities at vertices
stage
7
8
9
10A
(12)
(12)
(8)
(11)
**
*
C
ena
gRNA1
gRNA2
1 kB
ena1.8
Ena (isoform F)
EVH1
PRR
EVH2
5G2
100 aa
D
Act5c>GFP::FP4-mito clones (GFP +)
stage 8
E-Cad
GFP::FP4-mito
Ena
F-actin
E
ena1.8 clones (GFP -)
ena1.8
GFP (wt)
E-Cad
Ena
GFP
F-actin
F
Act5c>ena clones (RFP +)
Act5c>ena
E-Cad
RFP (clone)
Ena
F-actin
G
stage 8
F-actin intensities at vertices
(12)
(10)
(11)
control
Act5c> FP4-mito
control
ena1.8
control
Act5C> ena
H
ena1.8 clones (GFP -)
stage 8
stage 10A
GFP (wt)
Dextran
I
stage 8
stage 10A
Patency Index (%)
(10)
(9)
control
ena1.8
control
ena1.8

**Figure S4 (related to Figure 4)**

**Enabled/VASP is dispensable for vertex-associated F-actin formation in follicle cells.**

**(A)** Apical sections of FC vertices in living follicles expressing E-Cad::RFP (magenta) and GFP::Ena (green). Close-up views show single vertices marked by rectangles in overview. Single channel shows GFP::Ena intensities color-coded using heatmap. Note that GFP::Ena accumulates near vertices at stage 7 and 8 and declines by stage 9.

**(B)** Quantification of GFP::Ena signals (normalized) at vertices. Number of follicles (n) analyzed is indicated. P-values (Pairwise Wilcoxon Rank Sum Test): * p ≤ 0.05, ** p ≤ 0.01, *** p ≤ 0.001, **** p ≤ 0.0001.

**(C)** Schematic representation of the *ena*. locus and of Ena protein isoform F. *ena*$^{1.8}$ contains a deletion of 5328 bp and an insertion of 305 bp non-coding DNA at the breakpoint, resulting in premature stop codons in all annotated *ena* isoforms. The deletion removes most of the *ena* coding sequence, including the EVH1 (Ena-VASP-homology-1) domain, central Proline-rich regions (PRR) and part of the EVH2 (Ena-VASP-homology-2) domain. The region containing the epitope detected by the anti-Ena monoclonal antibody (5G2) is indicated.

**(D)** Apical sections of FC vertices in stage 8 follicles expressing GFP::FP4-mito (yellow) in clones, stained for E-Cad (magenta) and Ena (cyan; top) or E-Cad and F-actin (Phalloidin, cyan; bottom). Single channels show Ena or F-actin intensities color-coded using heatmap. Close-up views show single vertices marked by rectangles in overview.

**(E)** Apical sections of FC vertices in fixed stage 8 follicles carrying *ena*$^{1.8}$ homozygous clones (marked by absence of GFP, yellow), stained for E-Cad (magenta) and Ena (cyan; top) or E-Cad and F-actin (Phalloidin, cyan; bottom). Single channels show Ena or F-actin intensities color-coded using heatmap.

**(F)** Apical sections of FC vertices in fixed stage 8 follicles overexpressing Ena in clones (marked by RFP-nls, yellow), stained for E-Cad (magenta) and Ena (cyan; top) or E-Cad and F-actin (Phalloidin, cyan; bottom). Single channels show Ena or F-actin intensities color-coded using heatmap.

**(G)** Quantification of F-actin signals (normalized) at vertices in control cells and in clones either overexpressing Ena or expressing FP4-mito, and in *ena*$^{1.8}$ mutant clones (all stage 8). P-values (Welch's two-sample t-test): * p ≤ 0.05, ** p ≤ 0.01, *** p ≤ 0.001, **** p ≤ 0.0001.

**(H)** Basolateral sections of follicles carrying *ena*$^{1.8}$ homozygous clones (marked by absence of GFP, green). Dextran (magenta) marks intercellular spaces. Note that intercellular spaces at stage 10A are not altered upon loss of Ena in *ena*$^{1.8}$ clones.

**(I)** Quantification of patency index *ena*$^{1.8}$ clones and control cells. Number of follicles (n) analyzed is indicated. P-values (Welch's two-sample t-test): * p ≤ 0.05, ** p ≤ 0.01, *** p ≤ 0.001, **** p ≤ 0.0001.

Scale bars: (A-C, F) 5 µm, close-ups 1 µm, (H) 10 µm.

### Figure S5

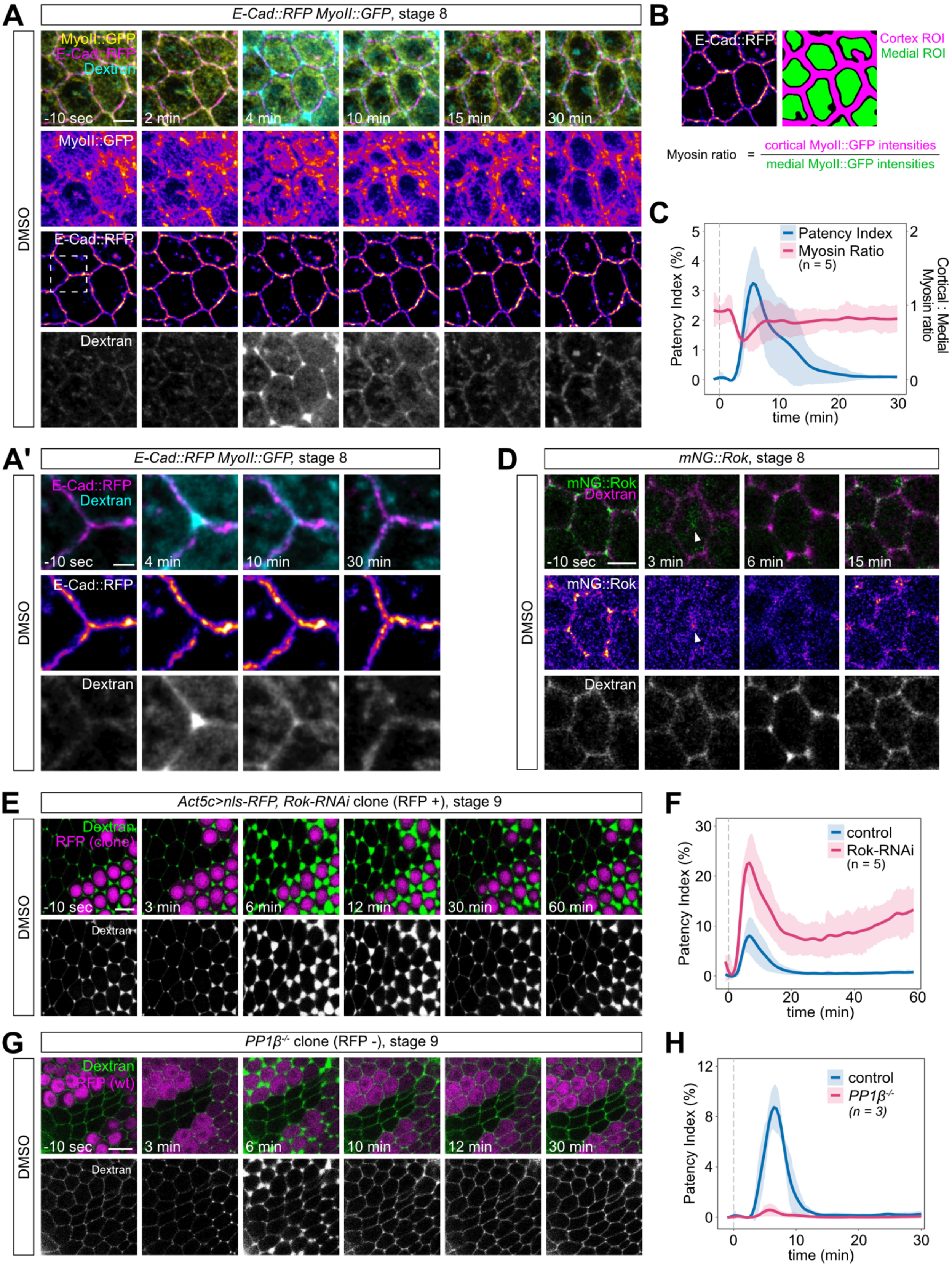
A
E-Cad::RFP MyoII::GFP, stage 8
DMSO
MyoII::GFP
E-Cad::RFP
Dextran
-10 sec
2 min
4 min
10 min
15 min
30 min
B
E-Cad::RFP
Cortex ROI
Medial ROI
Myosin ratio = cortical MyoII::GFP intensities / medial MyoII::GFP intensities
C
Patency Index
Myosin Ratio
(n = 5)
Patency Index (%)
Cortical : Medial Myosin ratio
time (min)
A'
E-Cad::RFP MyoII::GFP, stage 8
D
mNG::Rok, stage 8
mNG::Rok
3 min
6 min
E
Act5c>nls-RFP, Rok-RNAi clone (RFP +), stage 9
RFP (clone)
12 min
60 min
F
control
Rok-RNAi
G
PP1β-/- clone (RFP -), stage 9
RFP (wt)
H
PP1β-/-
(n = 3)

**Figure S5 (related to Figure 5)**

**Dynamics of E-Cad, Myosin II, and Rok in main body follicle cells after hypertonic shock**

**(A)** Stills (z-projections of apical sections) from pre-patent (stage 8) follicle expressing MyoII::GFP (yellow) and E-Cad::RFP (magenta) exposed to DMSO-induced hypertonic shock at t=0 min. Single channels show MyoII::GFP, E-Cad::RFP (heatmap) and dextran (grey). Dashed rectangle indicates position of closeups shown in (A'). Images were deconvolved using Leica Lightning Deconvolution. See also Video S5.

**(A')** Close-ups from (A). Merge shows E-Cad::RFP (magenta) and dextran (cyan), single channels show E-Cad::RFP (heatmap) and dextran (grey). See also Video S4.

**(B)** E-Cad::RFP (heatmap) and segmentation of cortical (magenta) and medial (green) regions based on E-Cad signal. The ratio of cortical : medial MyoII::GFP mean intensities ("Myosin ratio") was calculated and plotted in (C).

**(C)** Quantification of patency index and Myosin ratio (normalized) over time. Mean values and standard deviations, smoothed using a running median, are shown. Sample size (n) is indicated.

**(D)** Stills (apical sections) of pre-patent (stage 8) follicle expressing mNG::Rok (green) exposed to DMSO-induced hypertonic shock at t=0 min. Single channels show mNG::Rok (heatmap) and dextran (grey). Images were deconvolved using Leica Lightning Deconvolution. Arrowhead indicates medial accumulation of mNG::Rok after osmotic shock. See also Video S6.

**(E)** Stills (basolateral sections) of early patent (stage 9) follicle expressing *Rok* dsRNA in clone (marked by RFP-nls, magenta) exposed to DMSO-induced hypertonic shock at t=0 min. Single channel shows dextran (grey). Note that intercellular gaps in Rok-depleted tissue are enlarged and remain open when gaps in wild-type tissue close. See also Video S7.

**(F)** Quantification of patency index over time in control cells and in *Rok*-dsRNA-expressing cells. Mean values and standard deviations, smoothed using a running median, are shown. Sample size (n) is indicated.

**(G)** Stills (basolateral sections) of early patent (stage 9) follicle carrying *PP1β* (*flw*$^{FP41}$) clone (marked by absence of RFP, magenta) exposed to hypertonic shock (DMSO) at t=0 min. Single channel shows dextran (grey). Note that intercellular gaps in *PP1β* mutant tissue do not open comparable to surrounding wild-type tissue. See also Video S8.

**(H)** Quantification of patency index over time in control cells and in *PP1β* clone. Mean values and standard deviations, smoothed using a running median, are shown. Sample size (n) is indicated.

Scale bars: (A, D) 5 µm, (A') 2 µm, (E, G) 10 µm.

#### Supplemental Videos

##### Video S1 (related to Figure 1 and Figure S2)

**Laser ablations of apical FC vertices.**

Time-lapse movies of follicles expressing E-Cad::GFP (greyscale). Stages and time (sec) are indicated. Single vertices were ablated at t=0 sec. Note that neighboring vertices are displaced after laser ablation in stage 7 and 8 follicles, whereas little displacement is observed after laser ablation in stage 9 follicle. See also Fig. S2.
Scale bar: 5 µm.

##### Video S2 (related to Figure 4 and Figure 5)

**Osmotic treatment of cultured follicles.**
Time-lapse movies (basolateral sections) of stage 9 follicles stained with CellMask (magenta) and incubated in fluorescent dextran (green). Follicles were exposed to osmotic shocks with the indicated reagent at t=0 min. Note that addition of PBS or Sucrose induces more rapid vertex opening than addition of DMSO. Hypotonic treatment with water does not cause vertex opening. Time (min) is indicated. See also Fig. 5A.
Scale bar: 5 µm.

##### Video S3 (related to Figure 4 and Figure 5)

**Hypertonic treatment of stage 5 follicles containing interphase and mitotic cells.**
Time-lapse movies (basolateral sections) of FCs in interphase (left) or mitosis (right) in stage 5 follicles incubated with CellMask (magenta) and fluorescent dextran (green). Follicles were exposed to hypertonic shock with DMSO at t=0 min. Note that junctions of the dividing cell and at the cleavage furrow show transient leaks. Time (min) is indicated. See also Fig. 5E.
Scale bar: 5 µm.

##### Video S4 (related to Figure 5)

**Hypertonic treatment of follicle expressing E-Cad::RFP.**
Closeup of single vertex from Video S5. Time-lapse movie (average projection of apical sections) of stage 8 follicle expressing E-Cad::RFP (magenta). Fluorescent dextran (green) marks intercellular spaces. Single channels show E-Cad::RFP and dextran in grey. The follicle was exposed to hypertonic shock with DMSO at t=0 min. Time (min) is indicated. Images were deconvolved using Leica Lightning Deconvolution. See also Fig. S5A'.
Scale bar: 2 µm.

##### Video S5 (related to Figure 5)

**Hypertonic treatment of follicle expressing MyoII::GFP and E-Cad::RFP.**
Time-lapse movie (average projection of apical sections) of stage 8 follicle expressing MyoII::GFP (yellow) and E-Cad::RFP (magenta). Fluorescent dextran (cyan) marks intercellular spaces. Single channels show MyoII::GFP (heatmap) and dextran (grey). The follicle was exposed to hypertonic shock with DMSO at t=0 min. Time (min) is indicated. Images were deconvolved using Leica Lightning Deconvolution. See also Fig. S5A.
Scale bar: 5 µm.

##### Video S6 (related to Figure 5)

**Hypertonic treatment of follicle expressing mNG::Rok.**
Time-lapse movie (maximum projection of apical sections) of stage 8 follicle expressing mNG::Rok (green). Fluorescent dextran (magenta) marks intercellular spaces. Single channels show mNG::Rok (heatmap) and dextran (grey). The follicle was exposed to hypertonic shock with DMSO at t=0 min. Time (min) is indicated. Images were deconvolved using Leica Lightning Deconvolution. See also Fig. S5D.
Scale bar: 5 µm.

**Video S7 (related to Figure 5)**

**Hypertonic treatment of mosaic follicle carrying Rok-depleted clone.**
Time-lapse movie (basolateral section) of stage 9 follicle expressing *Rok* dsRNA in mRFP-nls-marked clone (magenta). Fluorescent dextran (green) marks intercellular spaces. Single channel shows dextran (grey). Follicle was exposed to hypertonic shock with DMSO at t=0 min. Note that vertices in Rok-depleted tissue fail to close after osmotic shock-induced opening. See also Fig. S5E.
Scale bar: 10 µm.

**Video S8 (related to Figure 5)**

**Hypertonic treatment of mosaic follicle carrying *PP1β* mutant clone.**
Time-lapse movie (basolateral section) of stage 9 follicle carrying *PP1β* mutant clone marked by absence of mRFP-nls (magenta). Fluorescent dextran (green) marks intercellular spaces. Single channel shows dextran (grey). Follicle was exposed to hypertonic shock with DMSO at t=0 min. Note that vertices in control tissue open, while vertices in *PP1β* clone remain closed after hypertonic shock. Time (min) is indicated. See also Fig. S5G.
Scale bar: 10 µm.

**Video S9 (related to Figure 4 and Figure 5)**

**Hypertonic treatment of follicle expressing Lifeact::GFP.**
Time-lapse movie (apical section) of stage 9 follicle expressing Lifeact::GFP (green) driven by CY2-Gal4. Fluorescent dextran (magenta) marks intercellular spaces. Single channel shows Lifeact::GFP intensities (heatmap). Follicle was exposed to hypertonic shock with DMSO at t=0 min. Note that Lifeact::GFP signals decrease before vertices open and reappear at closing vertices. Time (min) is indicated. See also Fig. 5G.
Scale bar: 5 µm.

**Video S10 (related to Figure 4 and Figure 5)**

**Hypertonic treatment of follicle expressing Lifeact::GFP treated with Latrunculin A.**
Time-lapse movie (apical section) of stage 9 follicle expressing Lifeact::GFP (green) driven by CY2-Gal4. Fluorescent dextran (magenta) marks intercellular spaces. Single channel shows Lifeact::GFP intensities (heatmap). The follicle was treated with Latrunculin A (2 µM) before imaging and exposed to hypertonic shock with DMSO at t=0 min. Note that intercellular gaps are present before hypertonic shock and that gaps fail to close . Time (min) is indicated. See also Fig. 5I.
Scale bar: 5 µm.

**Video S11 (related to Figure 6)**

**Optogenetic activation of Rho signaling followed by hypertonic treatment.**

Time-lapse movie (basolateral section) of stage 9 follicle expressing CIBN::pmGFP (yellow) and CRY2-RhoGEF::mCherry (magenta, grey in middle panel) in clone. Fluorescent dextran (cyan, grey in right panel) marks intercellular spaces. Rho signaling was activated by illumination with blue light at t=0 min and hypertonic shock with DMSO was applied at t=10 min. Note that vertices do not open in tissue with activated Rho, while vertices in neighboring control cells open after hypertonic shock. Time (min) is indicated. See also Fig. 6A.
Scale bar: 10 µm.

**Table S1:** ***Drosophila*** **genotypes**

| Figure | Panel | Genotypes |
|---|---|---|
| 1 | A, B | *; Fs(2)Ket-GFP / CyO; ubi-PH-RFP* |
| | C, D | *w; E-Cad::3xtagRFP / MyoII::3xGFP; +; +* |
| | E | *w; E-Cad::3xtagRFP / Vinc::GFP; +; +* |
| 2 | A | *mNeonGreen::Rok / w; E-Cad::3xtagRFP / +; +; +* |
| | B | *w; E-Cad::3xGFP; +; +* |
| | C, D | *y w Act5c>CD2>Gal4 / y w hs-Flp*$^{122}$ *; UAS-mCherry-nls / +; UAS-Rok-RNAi (JF03225) /+* |
| 3 | B,F,G | *ubi-mRFP-nls hs-Flp FRT19A / flw*$^{FP41}$ *g*$^{2}$ *f*$^{1}$ *FRT19A; +; +; +* |
| | I,K | *y w Act5c>CD2>Gal4 / w; UAS-mCherry-nls / UAS-MHC*$^{DN}$*::YFP; hs-Flp / +; +* |
| 4 | A | *w / y w; E-Cad::3xtagRFP / Tj-Gal4; UAS-Lifeact::mBaoJin / +* |
| | C, D | *w; E-Cad::3xGFP; +; +* |
| | F, H | *y w hs-Flp*$^{122}$ */ w; +; FRT82B ubi-mRFP-nls / FRT82B ssh*$^{1-11}$ |
| 5 | A,C,E,G | *w; CY2-Gal4 UAS-lifeact::GFP; +; +* |
| 6 | B | *y w Act5c>CD2>Gal4 / y w hs-Flp*$^{122}$*; UASp-CIBN::pmGFP / +; UASp-RhoGEF-CRY2-mCherry / +; +* |
| | D | *y w Act5c>CD2>Gal4 / y w hs-Flp*$^{122}$*; +; UASp-RhoGEF-CRY2-mCherry / +; +* |
| | | *y w Act5c>CD2>Gal4 / y w hs-Flp*$^{122}$*; UASp-CIBN::pmGFP / +; UASp-RhoGEF-CRY2-mCherry / +; +* |
| S1 | A | *w / y w; E-Cad::3xtagRFP / Tj-Gal4; UAS-Lifeact::mBaoJin / +* |
| | B | *w; E-Cad::3xtagRFP / UAS-Lifeact-Halo2; GR1-Gal4 / +* |
| | C | *w; E-Cad::3xtagRFP / + ; GFP::Jub / +; +* |
| | D | *w; E-Cad::3xtagRFP / + ; +; Zyx::GFSTF / +* |
| S2 | B | *w; E-Cad::3xGFP; +; +* |
| S3 | A, C | *w*$^{1118}$ |
| | | *w, LIMK1*$^{2}$ |
| S4 | A | *w; E-Cad::3xtagRFP / +; ubi-GFP::Ena / +; +* |
| | D | *y w Act5c>CD2>Gal4 / y w hs-Flp*$^{122}$*; +; UAS-FP4-mito-GFP / +; +* |
| | E, H | *y w / y w hs-Flp*$^{122}$*; FRTG13 ubi-nls-GFP / FRTG13 ena*$^{1.8}$*; +; +* |
| | F | *y w Act5c>CD2>Gal4 / y w hs-Flp*$^{122}$*; UAS-mCherry-nls / +; UAS-ena-wt / +; +* |
| S5 | A,A',B | *w; E-Cad::3xtagRFP ; MyoII::3xGFP; +* |
| | D | *mNeonGreen::Rok; + ; + ; +* |
| | E | *y w Act5c>CD2>Gal4 / y w hs-Flp*$^{122}$ *; UAS-mCherry-nls / +; UAS-Rok-RNAi (JF03225) /+* |
| | G | *ubi-mRFP-nls hs-Flp FRT19A / flw*$^{FP41}$ *g*$^{2}$ *f*$^{1}$ *FRT19A; +; +; +* |
